# KLF15 regulates sulfur amino acid metabolism through Cystathionine gamma-lyase

**DOI:** 10.64898/2026.08.28.746943

**Authors:** Zahra Mehrazad Saber, Yoshinori Takeuchi, Samia Karkoutly, Masato Higaki, Tsolmon Mendsaikhan, Rika Saikawa, Yuichi Aita, Yuki Murayama, Akito Shikama, Yukari Masuda, Naoya Yahagi

## Abstract

High-protein diets increase hepatic sulfur amino acid metabolism, but the underlying transcriptional mechanisms remain unclear. This study investigated whether Krüppel-like factor 15 (KLF15) directly regulates cystathionine γ-lyase (CTH), a key enzyme linking methionine transsulfuration to hydrogen sulfide (H₂S) and taurine production. Promoter-reporter assays, electrophoretic mobility shift assays, and chromatin immunoprecipitation identified two functional KLF15-binding elements, designated 1-1 and 2-2, within the proximal *Cth* promoter. Mutation of either element attenuated KLF15-dependent promoter activation, whereas mutation of both largely abolished it. In vivo luciferase imaging further demonstrated that these elements were required for the hepatic transcriptional response to a high-protein diet. KLF15 loss of function reduced high-protein-diet-induced *Cth* expression and altered the hepatic sulfur amino acid profile. Methionine, cystathionine, and cystine accumulated, whereas taurine production and the high-protein-diet-induced increase in hepatic H₂S were attenuated. Gene expression analyses further indicated that KLF15 selectively regulates components of methionine, taurine, and H₂S metabolism rather than controlling the entire sulfur metabolic program. Collectively, these findings establish the high-protein diet-KLF15-CTH axis as a physiologically relevant transcriptional pathway that amplifies hepatic sulfur amino acid disposal and directs sulfur toward H₂S and taurine production.

**Graphical abstract:** 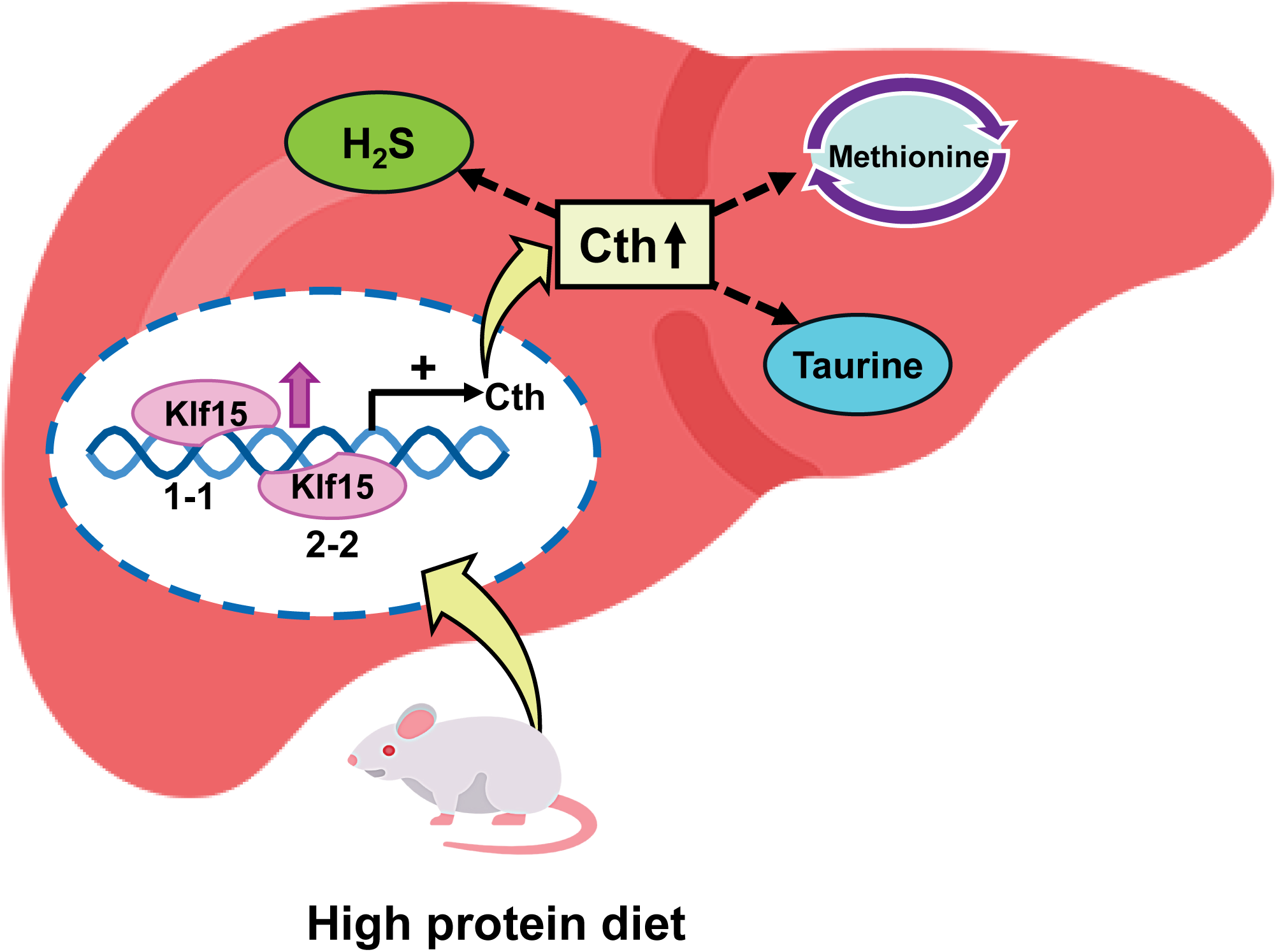

## Introduction

Dietary protein holds a unique regulatory position. Among the three macronutrients, protein has the greatest satiating effect per unit of energy, highest diet-induced thermogenesis, and is said to help preserve fat-free mass and maintain fat-mass loss during energy restriction much more effectively than isocaloric high-carbohydrate regimens (1–3). Increasing the proportion of energy derived from protein ameliorated weight-loss maintenance (2,4), reduced postprandial glycemic excursions and alleviated glycated hemoglobin in type 2 diabetes (5). It also attenuated the age-related loss of skeletal muscle mass in clinical trials, as reviewed in (6). Consequently, high-protein diets (HPD) are among the most widespread approaches for the prevention and management of obesity as well as sarcopenia (1,2).

A coordinated hepatic adaptation underlies these systemic benefits. As the first-pass organ for portal amino acid delivery, liver must dispose of amino acids ingested in excess of anabolic requirements. Classical enzymological work determined that urea cycle enzyme activities and aminotransferase activities rise corresponding to protein intake (7,8). Moreover, recent transcriptomic analyses have shown that HPD increases the hepatic expression of genes involved in the catabolism of essentially all twenty amino acids, and that this response is transcriptionally driven (9). As cysteine and methionine intake scales up in proportion with total food protein, it is notable that an HPD inflicts a large sulfur load. It is relatively unknown how liver transcriptionally regulates its sulfur amino acid disposal system relative to protein intake and what are the physiological consequences of such changes.

Cystathionine γ-lyase (CTH, CSE or cystathionase) catalyzes the second step of the transsulfuration pathway. The transsulfuration pathway starts with homocysteine generated by excess methionine from methionine cycle. Homocysteine is condensed with serine by cystathionine β-synthase (CBS) to form cystathionine. CTH is generally the cysteine production’s rate-limiting enzyme that cleaves cystathionine to cysteine, α-ketobutyrate and ammonia and therefore provides the principal route by which the sulfur atom of methionine is committed to cysteine (10,11). This step is confined to the liver, kidney and a few other tissues hence making hepatic CTH the principal gatekeeper for the entry of dietary methionine sulfur into the cysteine pool (10–12). Cysteine at this stage serves as a branch point for generation of two important products through different paths; one, hydrogen sulfide (H₂S) through desulfuration, a reaction catalyzed by CTH itself, and also by Cystathionine beta-synthase (CBS), 3-mercaptopyruvate sulfurtransferase (MPST/3-MST) and other lesser involved genes (13–15). The other, taurine through oxidation by cysteine dioxygenase (CDO1) to cysteine sulfinate and onward decarboxylation by cysteine sulfinate decarboxylase (CSAD), with hypotaurine subsequently oxidised by cysteamine dioxygenase (ADO) and flavin-containing monooxygenase 1 (FMO1) (11,12). CTH therefore sits at the junction of two functionally important molecules of active therapeutic interest — H₂S production and taurine synthesis — and any transcriptional input that sets CTH function will propagate to both.

Even though H_2_S was once considered an environmental toxin, nowadays it is established as the third mammalian gasotransmitter alongside nitric oxide and carbon monoxide (16,17). Its functions include vasorelaxant, cytoprotective, anti-inflammatory and mitochondrial-bioenergetic actions (17,18). 2-aminoethanesulfonic acid or taurine is one of the most abundant amino acids in the brain and spinal cord, leukocytes, heart and muscle cells, the retina, and indeed almost every tissue throughout the body, yet it is not incorporated into protein (19). Antioxidant and anti-inflammatory activity, regulation of energy metabolism, cytoprotection through modulation of endoplasmic reticular stress, gene expression regulation, inhibitory neuromodulator activity, modulation of Ca^2+^ homeostasis, osmoregulation, conjugation of bile acids, and post-transcriptional modification of mitochondrial tRNAs required for efficient mitochondrial translation are among its functions (20).

Krüppel-like factor 15 (KLF15) is a zinc-finger transcription factor, which binds to GC-rich regions and CACCC elements and mediates activation and/or repression of transcription (21). Liver, kidney, skeletal muscle, heart and adipose tissue have high expression levels of KLF15 and hepatic expression of KLF15 is rapidly induced by fasting and glucocorticoid receptor activation (22–26). Functionally, KLF15 coordinates fasting response by driving gluconeogenesis and the transcription of amino acid catabolic enzymes, including alanine aminotransferase (*Gpt*/*Alt*) and branched-chain aminotransferase (*Bcat2*) (23,24). Also, it assists mitochondrial acetyl-CoA synthetase expression and acetate utilization (26) and controls lipid flux and exercise adaptation in skeletal muscle (27). The absence of *Klf15* induced cardiac hypertrophy and fasting hypoglycaemia (23,28). Jeyaraj and colleagues demonstrated that *Klf15* orchestrates circadian nitrogen homeostasis on hepatic amino acid catabolism and urea cycle gene expression and on plasma amino acid concentrations. Disruption of *Klf15* abolished this rhythm and impaired nitrogen disposal (24). Independently, our group showed that fasting-induced Klf15 forms a complex with LXR/RXR specifically on the *Srebf1* promoter, thereby suppressing SREBP-1c and downstream lipogenic gene expression, and enabling the rapid switching between lipogenesis and gluconeogenesis during the early stages of fasting (29). Using an *in vivo* adenovirus–luciferase (Ad-luc) promoter analysis combined with a genome-wide transcription factor screen (TFEL scan) (30), we identified FoxO1 and FoxO3a as direct upstream activators of the liver-specific *Klf15* transcript, establishing a FoxO-Klf15 axis that switches macronutrient flow under the control of insulin signaling (31). KLF15 therefore constitutes a plausible transcriptional interface between protein intake and hepatic amino acid disposal. More recently, FoxO transcription factors were shown to regulate the urea cycle gene *Ass1* independently of Klf15, indicating that the hepatic protein response is composed of parallel, partially separable transcriptional arms (32).

Most directly relevant to the present work, our group characterized the hepatic HPD response in mice and used *Klf15*-knockout animals together with *in vivo* Ad-luc analysis to partition HPD-responsive genes into Klf15-dependent and Klf15-independent groups (9). That study reported that HPD-induced *Cth* expression is Klf15-dependent, whereas induction of aspartate aminotransferase (*Ast*) occurs independently of Klf15. This finding placed *Cth* within the Klf15 regulon (9).

In this study, for the first time, we hypothesized that Klf15 directly transactivates *Cth* by binding defined cis-elements within its proximal promoter, and that this Klf15-Cth axis mediates the high-protein-diet-induced increase in hepatic H₂S and taurine production and sulfur amino acid disposal.

## Results

### Identification of Klf15 transcription factor-binding region of the *Cth* gene promoter

We previously identified *Cth* as a KLF15-dependent, high-protein-diet (HPD)-responsive gene on the basis of transcript measurements in *Klf15*-deficient mice, but the responsible *cis*-regulatory element was not defined (9). To localize it, we first inspected publicly available Klf15 chromatin immunoprecipitation–sequencing (ChIP-seq) data from adult mouse liver (accessions SRX10012505 and SRX10012506) (Figure S1A). A single, sharp KLF15 enrichment peak was present immediately upstream of the *Cth* transcription start site on chromosome 3 (mm9, ≈157,588,000-157,590,000 bp). This signal rapidly declined across the following 2000 bp window (Figure 1A; Supplementary Figure S1A). To functionally dissect the genomic fragment spanned by the Klf15 peak, the area was subdivided into two adjacent subfragments, designated F1 and F2 (Figure 1A).

**Figure 1.**
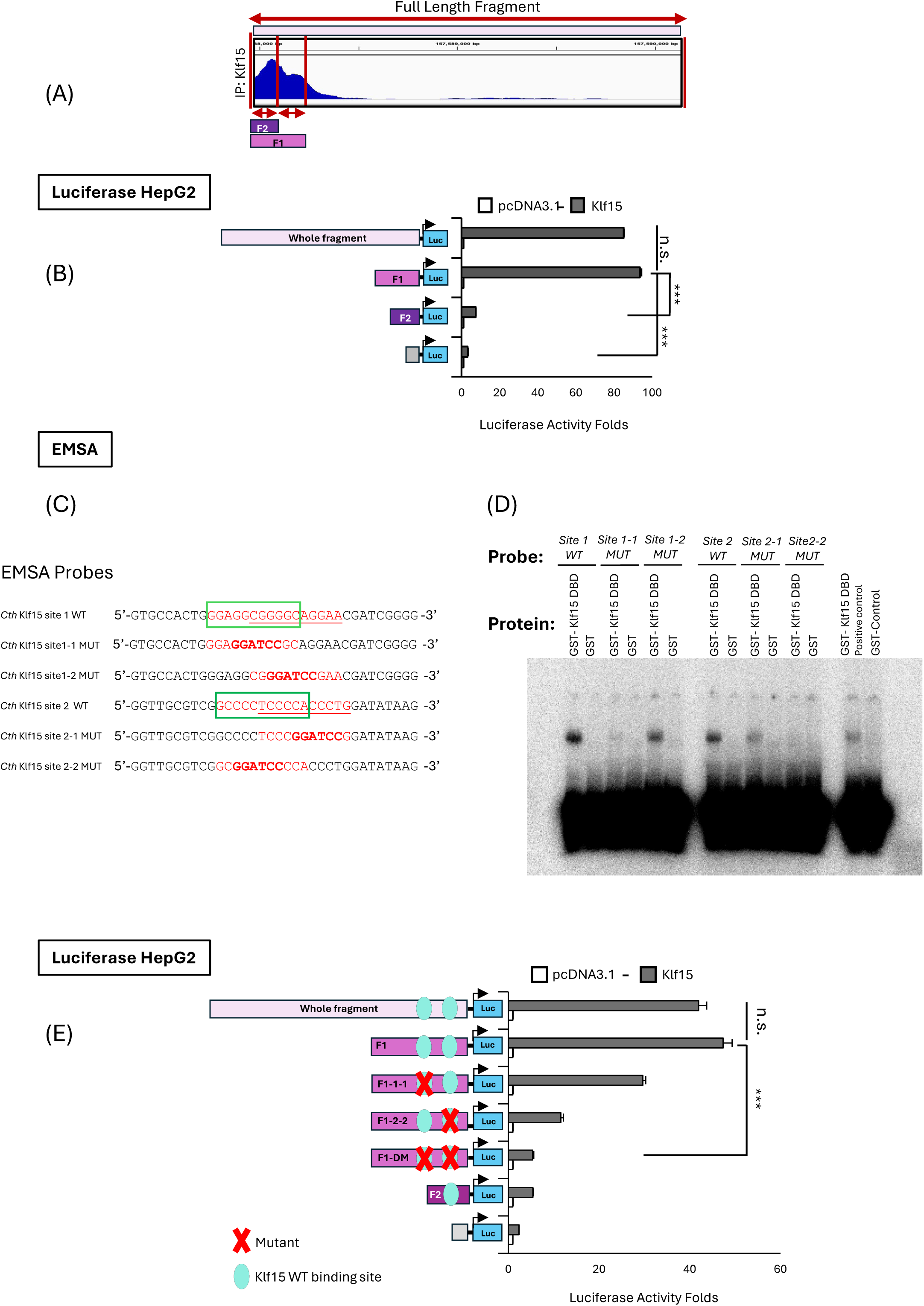
Identification of Klf15 transcription factor-binding region of the Cth gene promoter. **(A)** Top, the results of ChIP-seq with anti-Klf15 antibody in livers of mice obtained from ChIP-Atlas (ID: SRX10012506); bottom, simplified representation of the structure of the full fragment, fragment 1, fragment 2. (**B**) Promoter analysis of Klf15 on *Cth* gene and its fragments. Klf15 expression plasmid was co-transfected with the indicated *Cth* promoter firefly luciferase reporter plasmids in HepG2 cells. (**C** and **D**) Electrophoretic mobility shift assay (EMSA) using radiolabeled probe for (**C**) The four JASPAR predicted Klf15 binding sites in the fragment 1 of the *Cth* promoter in wild type shape. (**D**) the wild-type (WT) and mutant (Mut) predicted Klf15 binding site1 and site 2 incubated with GST and GST-Klf15 recombinant proteins. WT, wild-type; Mut, mutant. (**E**) analysis of Klf15 on mutant *Cth* promoter and its fragments. Klf15 expression plasmids were co-transfected with the indicated *Cth* promoter firefly luciferase reporter plasmids in HepG2 cells. Both (1–1) and (2–2) elements are functional, they contribute additively, and neither alone account for the full response; the 2-2 element makes the larger contribution in this cell-based assay. All values are presented as the mean with error bars representing the SEM. Datasets were assessed by Student’s *t*-test for unpaired samples. The differences were considered statistically significant if *P* < 0.05 (\**P* < 0.05 and \*\**P* <0.01).

To determine Klf15 transcription factor-binding sites in the *Cth* promoter region, we performed promoter analysis using F1 and F2 *Cth*-promoter-luc constructs in HepG2 cells. Possible Klf15 transcription factor binding sites in the promoter region of the *Cth* gene were identified using ChIP-Atlas (Figure S1C, S1D). F1 and F2 fragments along with the full-length fragment were cloned into luciferase reporter plasmid to be studied separately through evaluating the luciferase activity in HepG2 cells. As the data show, both full-length and F1 showed significant luciferase activity in response to Klf15 expression plasmids compared to F2 and to the control (Figure 1B).

To support these findings, an EMSA assay was performed. Therefore, six radiolabeled probes for the four high-scoring binding sites in fragment 1 based on JASPAR 2022 analysis results were designed (Figure S1C-D). These four binding sites were located in two pairs of overlapping binding sites. Within each cluster, two candidate elements were resolved and designated 1-1 and 1-2 (site 1 region) and 2-1 and 2-2 (site 2 region). Double-stranded probes corresponding to the wild-type (WT) sequences and to derivatives in which each individual element was replaced by a *Bam*HI linker (GGATCC) were synthesized (Figure 1C). EMSA results showed that the recombinant Klf15 protein binds to the DNA radiolabeled probe of both binding site number 1 (*Cth F1 site1 WT*) and binding site number 2 (*Cth F1 site2 WT*) (Figure 1C,D), while in each site, one of the mutated overlapping binding sites lost its activity and no band was observed. Klf15 therefore binds directly and selectively to elements 1-1 and 2-2, while the adjacent 1-2 and 2-1 elements are dispensable for binding under these conditions. (Figure 1D). GST alone did not produce a comparable shifted complex, confirming that the observed shifts were attributable to the KLF15 DNA-binding domain (Figure 1C,D).

Consistent with these findings, when we mutated these predicted binding sites in the fragment 1, the activation of the luciferase gene by Klf15 expression plasmids in HepG2 cells was completely abolished as shown in Figure 1E. Collectively, these findings identify the specific Klf15-binding site essential for promoter-mediated activation. The difference between F1 and F1 double-mutant (DM) was significant (***P < 0.001; Figure 1E), indicating that loss of both elements accounts for essentially all Klf15-dependent activity of the fragment, with element 2-2 contributing the larger share of the response in this assay.

### The identified region is required for hepatic *Cth* response to HPD

To determine whether this element responds to dietary protein *in vivo*, adenoviral reporter constructs were delivered to mouse liver by jugular-vein injection and hepatic luminescence was quantified by *in vivo* imaging (IVIS) in mice maintained on a low-protein diet (LPD) or a high-protein diet (HPD) (Figure 2) (29,33,34).

**Figure 2.**
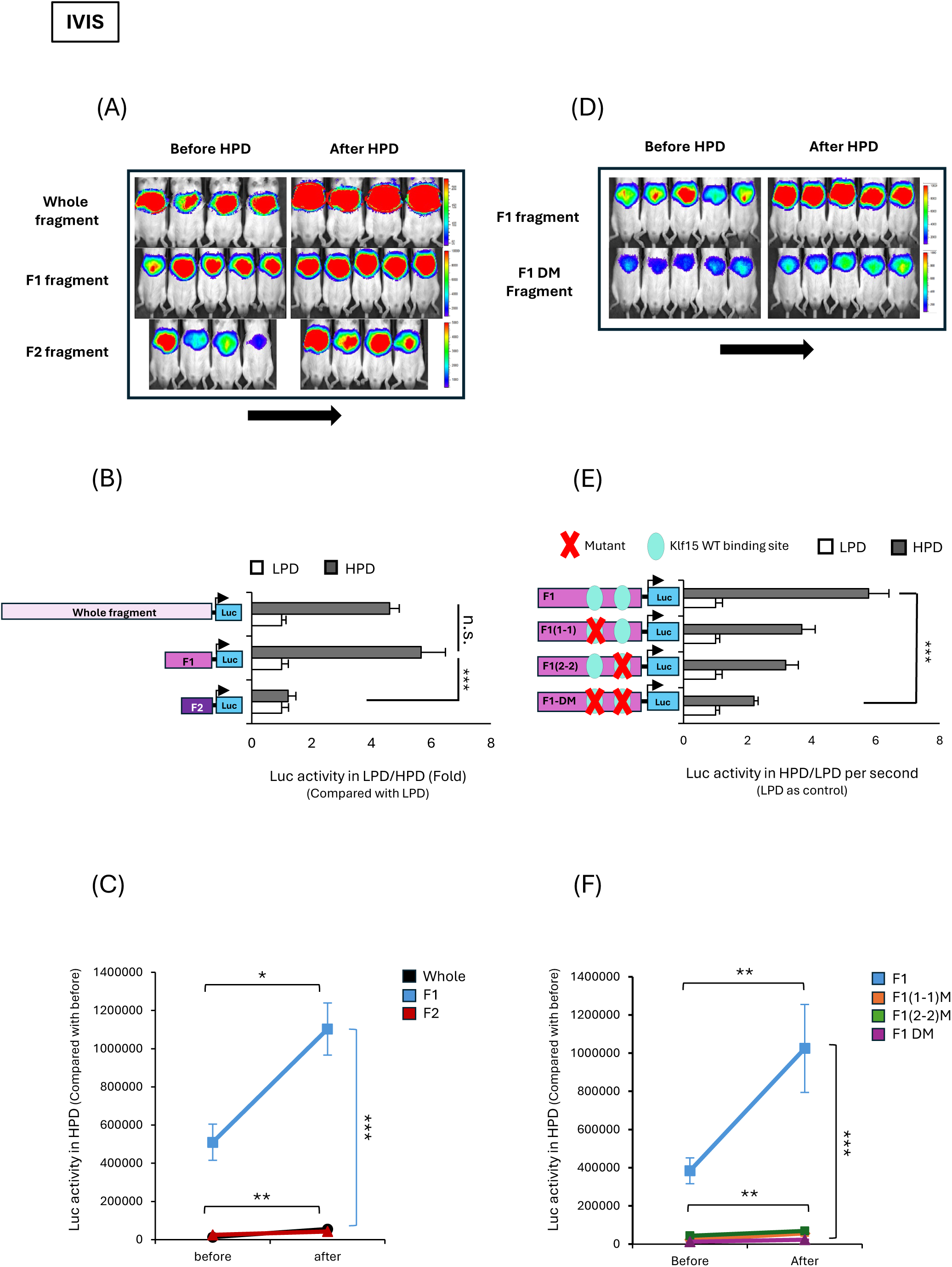
The identified region is required for hepatic *Cth* response to HPD. Luciferase reporters driven by the full-length *Cth* promoter fragment (Whole), the (F1) or (F2) sub-fragment, or F1 carrying mutated KLF15 elements — F1(1–1), F1(2-2), F1-DM — were delivered to the liver of ICR mice through jugular vein, which were then fed a low-(LPD) or high-protein diet (HPD) and imaged by IVIS. **(A)** Representative hepatic bioluminescence images for whole promoter, F1 and F2 fragments *in vivo* activities under HPD. **(B)** Hepatic luciferase activity under LPD or HPD, relative to the LPD group of each construct. F1’s reaction was significantly higher than F2. **(C)** Photon flux before and after transition to HPD. Post-HPD F1 exceeded both other constructs (***). **(D)** Representative images of mice fed HPD for wild-type fragments vs double mutated fragment. **(E)** Activity of F1 and its mutants relative to the corresponding LPD group. HPD responsiveness was progressively attenuated by mutation of either element and lowest in the double mutant. Cyan ovals, intact KLF15 sites; red crosses, mutated sites. **(F)** Photon flux before and after HPD for the constructs in (E). F1 response increased (**); all mutants remained low (***, F1 vs mutants after HPD). Data are mean ± SEM; *n* = 8 to 14 mice per group. Diet- and construct-dependent differences in (B) were assessed by the Kruskal-Wallis test with Bonferroni correction for multiple comparisons; group comparisons in (E) by one-way ANOVA with Tukey’s post hoc test; and paired before-versus-after measurements in (C) and (F) by the Wilcoxon signed-rank test. All analyses were performed with SPSS version 31.0.0 ; α = 0.05 (two-tailed). n.s., not significant; \**P* < 0.05; \*\**P* < 0.01; \*\*\**P* < 0.001.

We assessed the effect of HPD on transcriptional activity *in vivo* by engineering the promoter fragment containing the identified Klf15 binding sites (Figure 1) into a luciferase reporter plasmid. Then we compared them with the *Cth* native promoter (Figure 2B). The transcriptional activity was assessed by measuring luciferase activity with an IVIS imaging system before and after diet. As shown in Figure 2A-C the reporter activity of the native promoter and F1 fragment construct containing the wild-type Klf15 binding sites was significantly increased in the HPD-fed state compared to the LPD-fed state. On the other hand, F2 reporter activity was similar between HPD and LPD-fed states and significantly lower than those of F1 and whole promoter results. The same result was obtained using absolute photon counts in individual animals compared before and after the switch to HPD (Figure 2C). The F1 element is therefore both necessary and sufficient, among the fragments tested, to confer HPD responsiveness in the intact liver. The F1 reporter and its element mutants were next compared under LPD and HPD (Figure 2D-F).

Constructs containing the F1 fragment with mutated Klf15 binding sites (1–1) and (2–2) were made individually. An F1 fragment construct with double mutated binding sites was also generated. Results showed that mutation of either Klf15 element attenuated, and mutation of both largely eliminated, the transcriptional response of Cth promoter to dietary protein *in vivo* (Figure 2E). The same result was obtained using absolute photon counts in individual animals compared before and after the switch to HPD (Figure 2F).

### Klf15 occupies both *Cth* elements in hepatic chromatin under HPD

ChIP-qPCR was performed on liver chromatin from HPD-fed mice using an anti-Klf15 antibody or matched non-immune IgG. Both *Cth* elements were significantly enriched (both **P < 0.01), whereas little enrichment was observed with the IgG control or at negative control loci. The *Alt* promoter, a previously reported KLF15 target used here as a positive control locus (24), showed around 4.3-fold enrichment (**P < 0.01), whereas a gene-desert negative-control amplicon showed no enrichment (1.2-fold; n.s.) (Figure 3; primers in Supplementary Table S3). Klf15 therefore occupies both functionally required elements in native chromatin under high dietary protein (*n* = 6).

**Figure 3.**
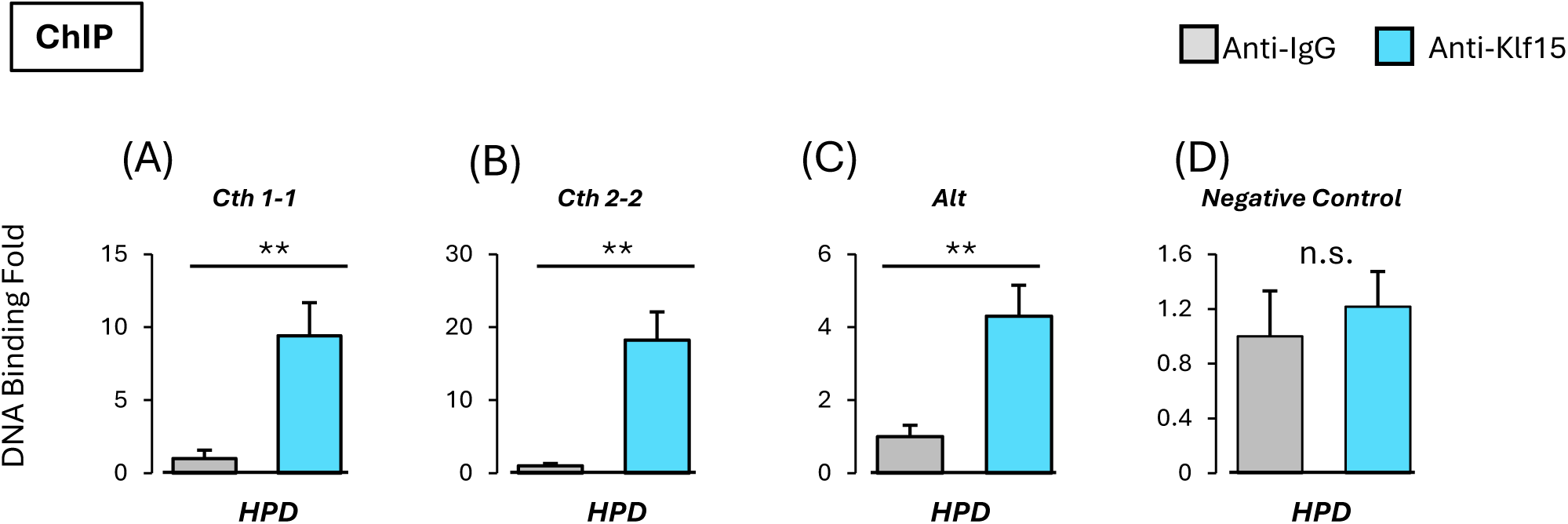
Klf15 occupies both *Cth* elements in hepatic chromatin under HPD. **(A-D)** Chromatin immunoprecipitation (ChIP) assay to determine the interaction of hepatic Klf15 proteins with *Cth* promoter region using anti-Klf15 antibody and normal mouse anti-IgG antibody as a control. (**A**, **B**) Klf15 proteins binding to *Cth* promoter in liver detected by using primer set designed around the identified Klf15 binding sites in Figure 1. (**C**) *Alt* promoter Klf15 transcription factor binding site was used as a positive control. (**D**) A gene-desert amplicon was used as negative control. All groups of mice were sacrificed in the light cycle, and liver samples were collected in the fed state following 3 days of ad libitum-fed HPD as indicated. Data were assessed using the unpaired two-tailed Student’s *t*-test. The differences were considered significant if *P* < 0.05 (\*\**P* < 0.01). Error bars mean SEM.

Taken together, this section and the previous one establish that KLF15 binds two defined elements in the proximal *Cth* promoter. These elements are occupied in liver chromatin *in vivo*, and they are required for both Klf15-driven and HPD-driven transactivation.

### Metabolic map linking dietary methionine to hepatic H₂S and taurine

The map shown in Figure 4 summarises the pathway through which hepatic sulfur amino acids are routed toward H₂S and taurine, and provides the interpretive framework for the metabolite measurements in Figure 5. Dietary methionine is activated to *S*-adenosylmethionine (SAM) by MAT and, via *S*-adenosylhomocysteine (SAH) and AHCY, converted to homocysteine, which is either remethylated back to methionine (MS/MTR; BHMT, using betaine derived from choline) or committed to the transsulfuration pathway (18). In transsulfuration, CBS condenses homocysteine with serine to form cystathionine, and CTH (cystathionine γ-lyase) cleaves cystathionine to cysteine; both enzymes additionally generate the gasotransmitter H₂S, and CTH can produce H₂S directly from cysteine and cystine (18). Cysteine constitutes the branch point: it is oxidised by CDO1 to cysteine sulfinate, decarboxylated by CSAD to hypotaurine, and converted to taurine by ADO/FMO1 (11). H₂S is generated convergently by CTH, CBS, the MPST(3-MST)/CAT couple, CARS2, and MTO (from methanethiol) (18). Within this scheme, CTH occupies a hub position — feeding H₂S directly while supplying the cysteine pool that drives taurine synthesis — such that a change in CTH activity is predicted to shift the balance between accumulation of upstream substrates (methionine, cystathionine) and formation of the downstream end-product taurine.

**Figure 4.**
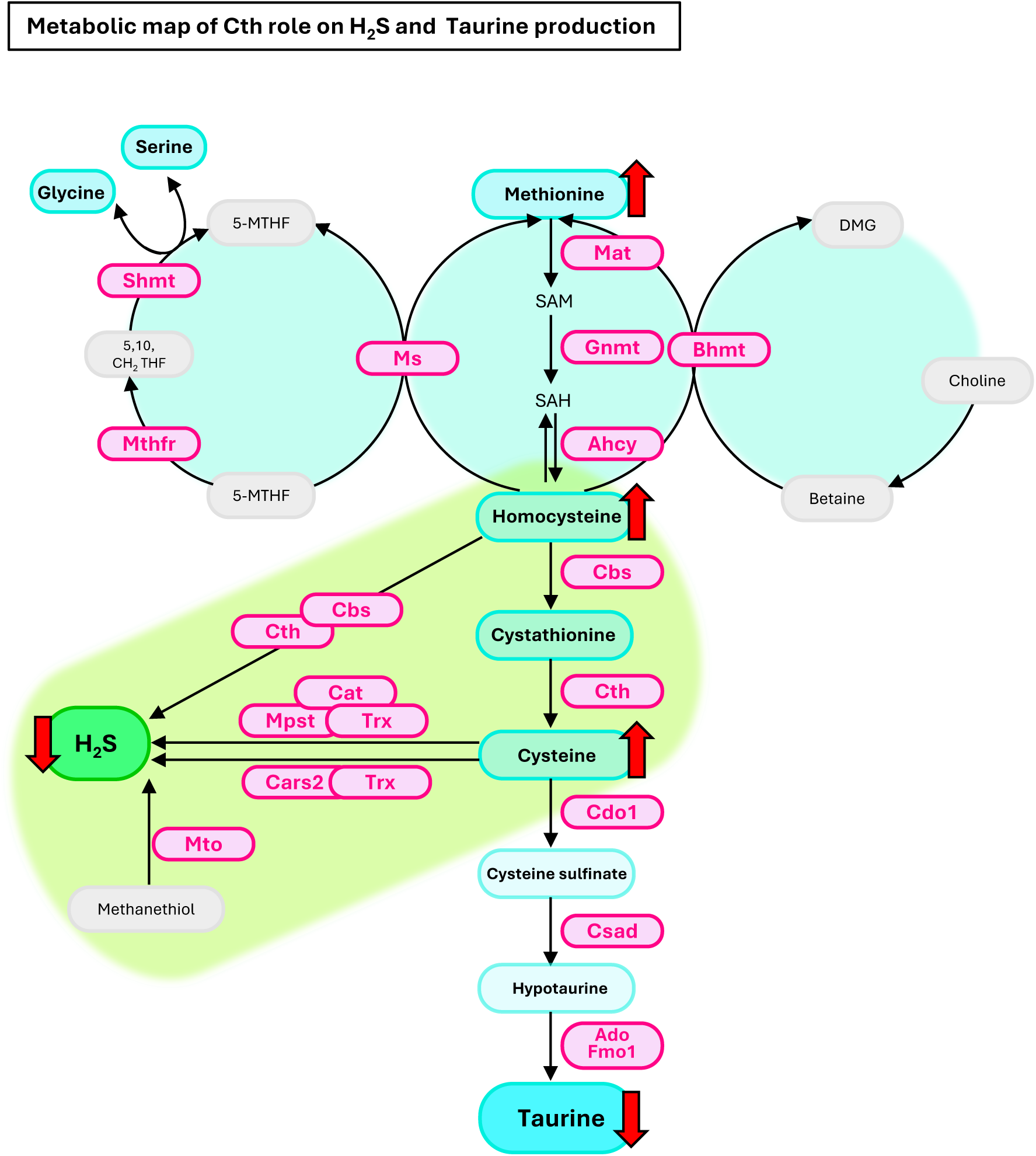
Metabolic map of the CTH-dependent route to hepatic H₂S and taurine production regulated by Klf15. Schematic of the methionine cycle, folate one-carbon metabolism, the transsulfuration pathway, and the cysteine→taurine branch. Metabolites are shown as nodes and enzymes as labelled steps (*Mat*, *Gnmt*, *Ahcy*, *Ms/Mtr*, *Bhmt*, *Shmt*, *Mthfr*, *Cbs*, *Cth*, *Mpst/3-Mst*, *Cat*, *Cars2*, *Mto*, *Cdo1*, *Csad*, *Ado*, *Fmo1*). CTH (cystathionine γ-lyase) occupies a hub position, generating H₂S directly and supplying cysteine for downstream taurine synthesis. The map provides the interpretive framework for the metabolite measurements in Figure 5.

**Figure 5.**
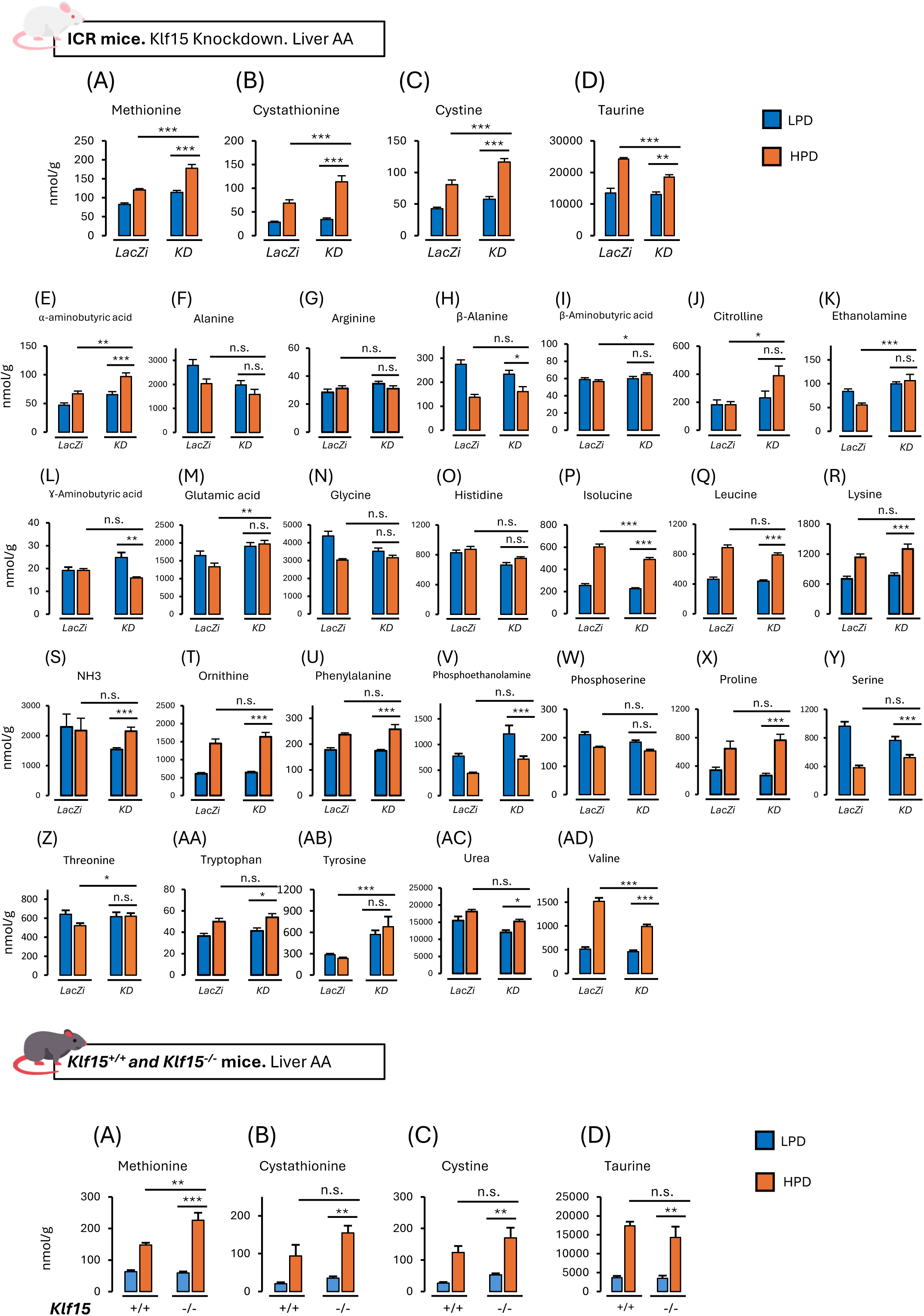
Acute hepatic *Klf15* knockdown alters the HPD-induced hepatic sulfur amino acid profile. Hepatic concentrations (nmol/g tissue) of free amino acids and related metabolites are shown under LPD (blue) and HPD (orange). **(A–AD)** Knockdown model: adenoviral *Klf15* knockdown in ICR mice (Ad-LacZi control vs. Ad-KLF15i, “KD”), *n* = 19 per group; panels A–D show the sulfur amino acids (methionine, cystathionine, cystine, taurine) and E–AD the remaining amino-acid panel. Separate panels A–D for the germline model: *Klf15*⁺ᐟ⁺ vs. *Klf15*⁻ᐟ⁻ mice, *n* = 6 per group (methionine, cystathionine, cystine, taurine). Data are mean ± SEM. Group comparisons were assessed by ANOVA with Tukey’s HSD post-hoc test for normally distributed data and by the Kruskal–Wallis test with Bonferroni correction for non-normally distributed data; \**P* < 0.05, \*\**P* < 0.01, \*\*\**P* < 0.001, n.s. = not significant.

### Acute hepatic *Klf15* knockdown alters the HPD-induced hepatic sulfur amino acid profile

To determine how KLF15 shapes the hepatic amino-acid response to protein loading, we quantified free amino acids and related metabolites in liver under LPD and HPD, in the acute knockdown (KD; ICR, panels A-AD) and constitutive knockout (KO; *Klf15*^-/-^, panels A-D) models (Figure 5). High-protein feeding broadly remodeled the hepatic free amino-acid pool in both control and Klf15-depleted livers, consistent with increased hepatic amino-acid catabolism under protein loading (9). The most pronounced and KLF15-sensitive changes, however, were concentrated in the transsulfuration–taurine axis mapped in Figure 4. The concentrations of 30 amino acids and related metabolites were measured in the livers of control LacZi-treated and KLF15-knockdown mice maintained on an LPD or HPD.

Transsulfuration intermediates including methionine, cystathionine and cystine were consistently elevated by HPD and further elevated by *Klf15* knockdown. Taurine rose steeply with HPD in both groups but reached a significantly lower level in knockdown livers. This suggests that KLF15 shapes the HPD-driven transsulfuration and taurine response (KD model, Fig. 5A–D). This substrate-up/product-down signature is the expected consequence of attenuated flux at the KLF15-regulated, CTH-centered steps, indicating that KLF15 sets the amplitude and downstream partitioning of the response rather than permitting or blocking it.

Aside from the three transsulfuration intermediates and taurine, HPD significantly increased isoleucine, leucine, valine, lysine, ornithine, phenylalanine, tyrosine, proline, NH₃ and urea, reflecting increased amino-acid oxidation and ureagenesis under protein loading (9). Among these targets, isoleucine and valine showed significant reduction in *Klf15* knockdown group in comparison with LacZi group. α-aminobutyric acid, citrulline and tyrosine showed significant increase in *Klf15* knockdown group in comparison with LacZi group. On the other hand, HPD decreased serine, phosphoethanolamine and β-alanine, significantly in both *Klf15* knockdown and LacZi groups. Ethanolamine fell with HPD in control livers but not in knockdown livers (n.s.); γ-aminobutyric acid fell with HPD only in knockdown livers. β-aminobutyric acid, glutamic acid and threonine showed slight reductions with HPD. This effect was removed in response to *Klf15* knockdown. Alanine, arginine, glycine, histidine and phosphoserine showed no significant diet or knockdown effect (n.s.). Together, these observations support a role for KLF15 in coordinating hepatic sulfur-containing amino acid handling during adaptation to increased dietary protein intake.

An equivalent panel was measured in constitutive knockouts (Figure 5). The transsulfuration signature was reproduced and amplified: methionine, cystathionine and cystine all rose with HPD and were higher in knockouts, although the genotype comparisons for cystathionine and cystine did not reach significance (n.s.). Taurine increased with HPD in both genotypes. In knockouts, although the pattern of lower taurine levels were visible, it also did not reach significance (n.s.) between genotypes. This blunting of the genotype effect in the constitutive model most plausibly reflects lifelong compensatory adaptation (e.g., upregulation of redundant transsulfuration/taurine routes) together with the systemic, whole-body nature of germline deletion, in contrast to the liver-restricted, acute loss achieved by adenoviral knockdown. Critically, the retention of significant HPD-induced increases in cystathionine, cystine and taurine in the complete absence of KLF15 reinforces that KLF15 is an amplitude modulator, not an obligatory gate, of this pathway.

### Hepatic H₂S accumulates on HPD in a Klf15-dependent manner

The transsulfuration pathway is the principal route of hepatic hydrogen sulfide (H₂S) generation, with cystathionine γ-lyase (CTH/CSE) contributing the majority of liver H₂S from cysteine (18). Previously, we had identified *Cth* as a KLF15-dependent, HPD-responsive target (9). We next asked whether KLF15 is required to translate dietary protein load into increased hepatic H₂S.

In control ICR mice, HPD significantly increased hepatic H₂S levels (Figure 6A). This establishes hepatic H₂S as a diet-responsive readout downstream of the sulfur amino-acid program. The H₂S increase was attenuated following hepatic *Klf15* knockdown, indicating that KLF15 is required for the full H₂S-producing response to an HPD (Figure 6B). We then examined constitutive germline loss of KLF15 in male mice (Figure 6C). In contrast to the knockdown, male *Klf15*^-/-^ liver retained a full HPD-induced H₂S response.

**Figure 6.**
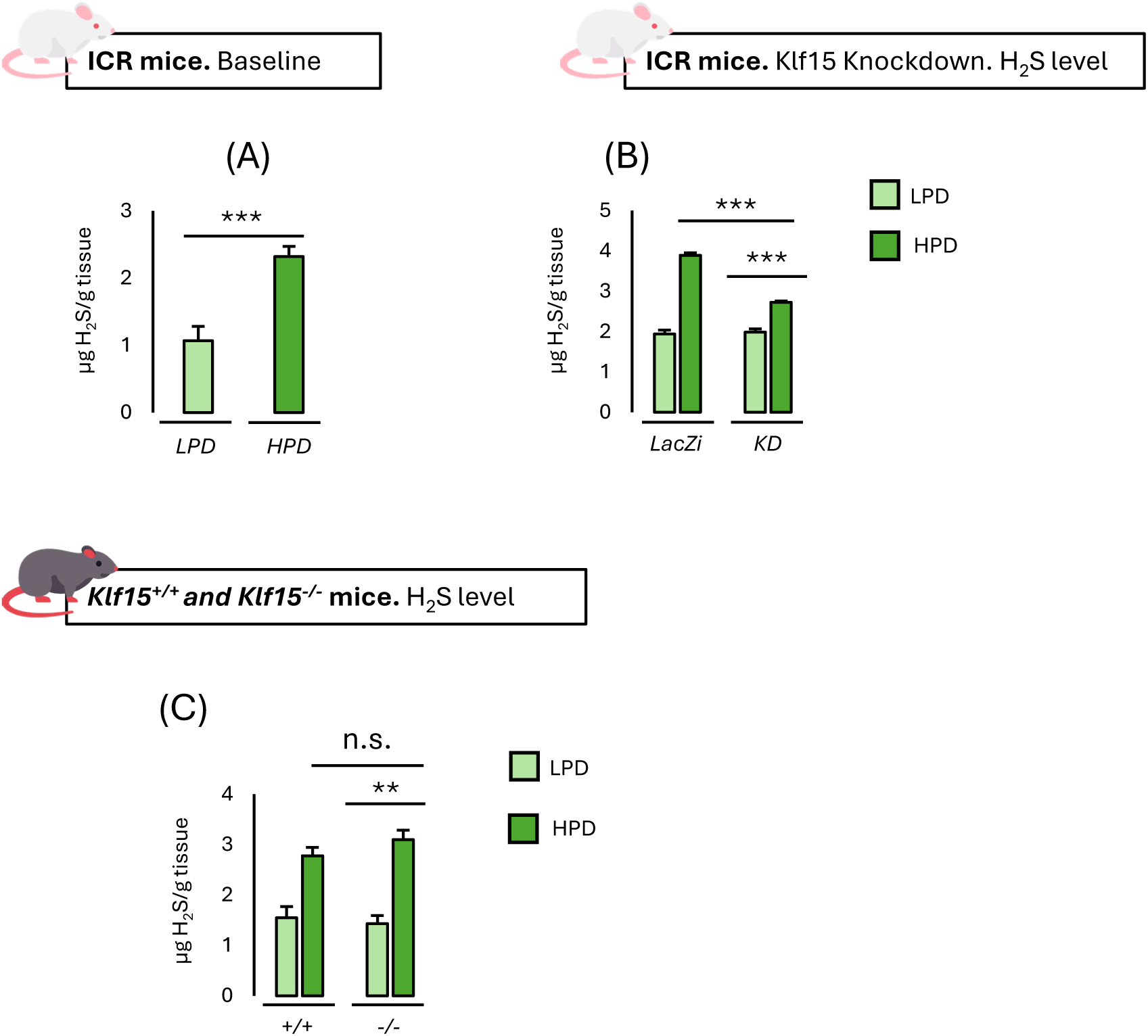
Hepatic H₂S Accumulates on HPD in a Klf15-Dependent Manner. All panels are from a single experiment using male mice. **(A)** Hepatic H₂S content (µg H₂S/g tissue) in male ICR mice fed a low-protein diet (LPD) or high-protein diet (HPD) (*n* = 6 per group). **(B)** Hepatic H₂S content in male ICR mice after adenoviral control (*LacZi*) or *Klf15* knockdown (*KD*), under LPD and HPD (*n* = 19 per group). **(C)** Hepatic H₂S content in male *Klf15*⁺ᐟ⁺ and *Klf15*⁻ᐟ⁻ mice under LPD and HPD (*n* = 6 per group). Data are mean ± SEM. Normally distributed data were analyzed by one-way ANOVA followed by Tukey’s HSD post hoc test; non-normally distributed data were analyzed by the Kruskal-Wallis test with Bonferroni correction for multiple comparisons. Distribution assessed by normality test Shapiro–Wilk. Analyses were performed in SPSS version 31.0.0 with α = 0.05. \**P* < 0.05, \*\**P* < 0.01, \*\*\**P* < 0.001; n.s., not significant. Absolute H₂S values are not directly comparable with Supplementary Figure S3 (independent experiments assayed with a refined H₂S protocol)

To determine whether the H₂S phenotype differs between sexes, we performed separate experiments in which females and males were analyzed independently (Supplementary Figure S3). In male mice, therefore, the two loss-of-function approaches diverge: In the knockdown model (Supplementary Figure S3A), the phenotype was consistent across sexes (F-*KD* vs M-*KD*, not significant). Acute KLF15 depletion thus blunts the HPD H₂S response equivalently in males and females. In the germline knockout model (Supplementary Figure S3B), by contrast, the effect of KLF15 loss was sexually dimorphic. In males, the HPD response was fully preserved. In females, the HPD response was retained but significantly dampened. Directly comparing knockout animals under HPD, males sustained higher hepatic H₂S than females. The male result is concordant with the male-only knockout experiment in Figure 6C (no significant *Klf15*^+/+^ vs *Klf15*^-/-^ difference under HPD), reproducing full preservation of the male response across two independent experiments.

### Methionine-cycle gene expression is selectively modulated by KLF15

By driving HPD-induced *Cth* (cystathionine γ-lyase), KLF15 promotes transsulfuration, the pathway that drains homocysteine — and thereby methionine-cycle carbon and sulfur — into cysteine synthesis. KLF15 is therefore positioned to limit hepatic methionine accumulation: when *Cth* induction is blunted, less homocysteine is committed to transsulfuration, which is compatible with the higher hepatic methionine observed after KLF15 loss. To characterize how the methionine cycle is reconfigured, hepatic transcript levels of methionine-cycle, remethylation and folate one-carbon genes were quantified by quantitative RT-PCR under LPD and HPD in the two KLF15 loss-of-function models: adenoviral *Klf15* knockdown in ICR mice and germline *Klf15*^-/-^ mice (Figure 7).

**Figure 7.**
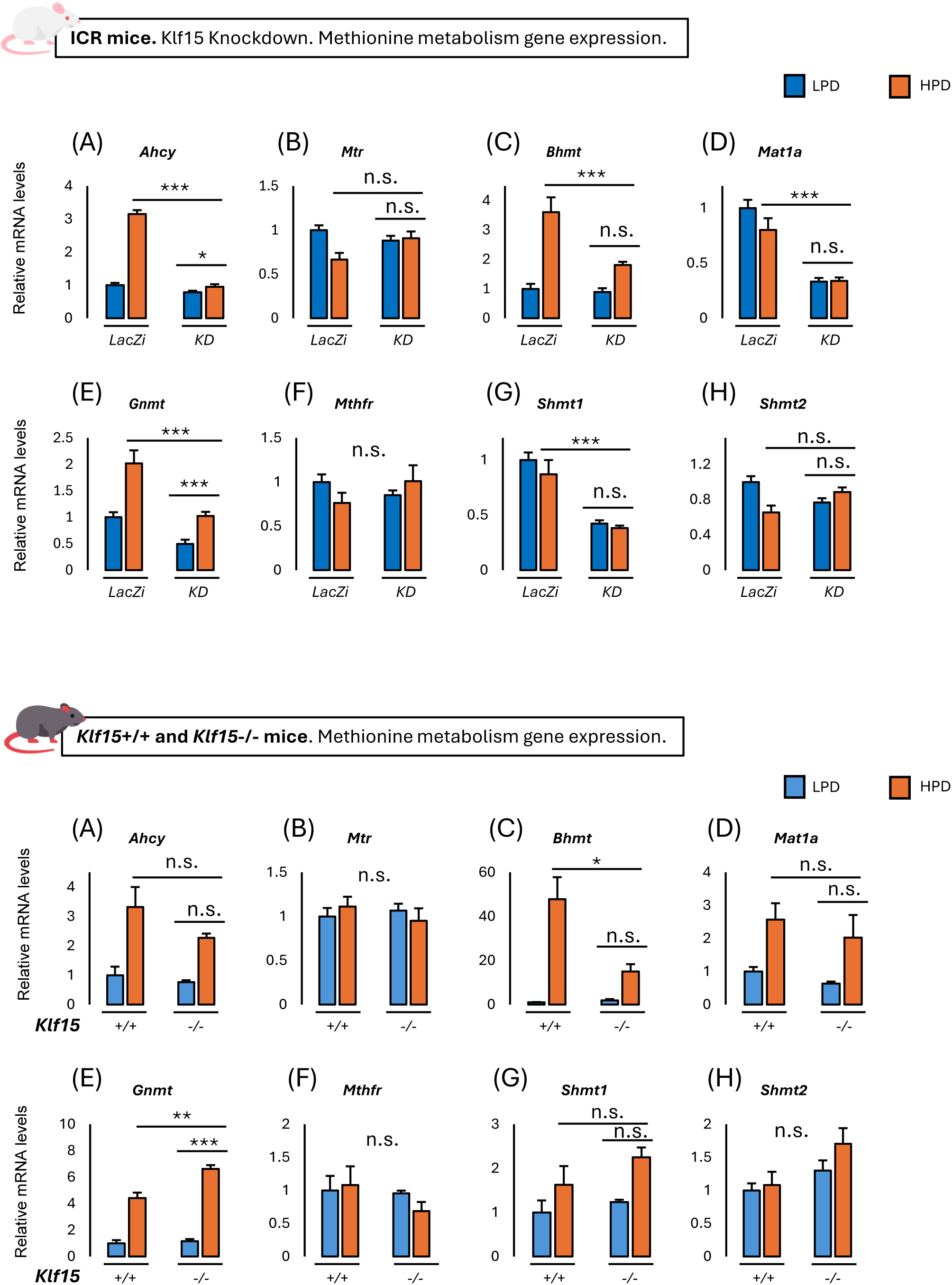
Methionine-cycle gene expression is selectively modulated by KLF15. Relative hepatic mRNA levels of methionine-cycle, remethylation and folate one-carbon genes measured by quantitative RT-PCR (normalized to *36B4*/*Rplp0*) in mice on low-protein diet (LPD, blue) or high-protein diet (HPD, orange). ICR mice injected with control (*LacZi*) or *Klf15*-knockdown (*KD*) adenovirus (Upper panels, A–H) (*n* = 10 per group). *Klf15*⁺/⁺ and *Klf15*⁻/⁻ mice (Lower panels, A–H) (*n* = 3 per group). Genes: **(A)** *Ahcy* (S-adenosylhomocysteine hydrolase); **(B)** *Mtr*/*Ms* (methionine synthase); **(C)** *Bhmt*(betaine-homocysteine S-methyltransferase); **(D)** *Mat1a* (methionine adenosyltransferase 1A); **(E)** *Gnmt* (glycine N-methyltransferase); **(F)** *Mthfr* (methylenetetrahydrofolate reductase); **(G)** *Shmt1* (cytosolic serine hydroxymethyltransferase); **(H)** *Shmt2* (mitochondrial serine hydroxymethyltransferase). Data are mean ± SEM (*n* = 3 to 10 per group). Groups were compared by one-way ANOVA with Tukey HSD post hoc test. \**P* < 0.05, \*\**P* < 0.01, \*\*\**P* < 0.001; n.s., not significant.

In the knockdown model, HPD induced *Ahcy*, *Bhmt* and *Gnmt* in control livers; each induction was significantly attenuated by *Klf15* knockdown and therefore showed a Klf15-dependent pattern. *Mat1a* and *Shmt1* were both reduced by *Klf15* knockdown and lost their diet responsiveness. *Mtr* was suppressed by HPD in controls only, *Shmt2* (mitochondrial isoform) differed modestly between groups and *Mthfr* was unchanged (n.s.).

In *Klf15*^-/-^ mice, *Bhmt* showed a large Klf15-dependent dietary response, and *Gnmt* was induced in both genotypes. *Ahcy* was induced by HPD in knockouts with no significant genotype difference (n.s.), *Shmt1*, *Mtr*, *Mat1a*, *Mthfr* and *Shmt2* (mitochondrial isoform) responses were unchanged (n.s.).

Several methionine-cycle genes (*Ahcy*, *Mat1a*, *Gnmt*, *Shmt1*) behaved differently between the acute knockdown and the constitutive knockout. Overall, HPD reconfigures the hepatic methionine cycle chiefly through induction of *Ahcy*, *Bhmt* and *Gnmt*, with KLF15 required for the full amplitude of these responses under acute knockdown. Together with the blunted *Cth*-mediated transsulfuration drainage documented in Figure 9, this remodeling is consistent with the greater hepatic methionine accumulation seen when KLF15 is reduced.

### Taurine-biosynthetic gene expression is reprogrammed by Klf15 loss

Because KLF15 drives HPD-induced *Cth* (cystathionine γ-lyase) and thereby transsulfuration flux, it is positioned to support hepatic taurine production indirectly by providing the supply of cysteine that feeds the dedicated taurine-biosynthetic pathway. We therefore asked whether the enzymes of taurine metabolism are themselves under KLF15 control, and whether their regulation accounts for the HPD-driven rise in hepatic taurine. Hepatic transcript levels of taurine-metabolism genes were quantified by quantitative RT-PCR under LPD and HPD in the two KLF15 loss-of-function models: adenoviral *Klf15* knockdown in ICR mice and germline *Klf15*⁻/⁻ mice. The genes assayed span the principal biosynthetic route (*Cdo1* → *Csad*), the alternative cysteamine route and hypotaurine oxidation (*Ado*, *Fmo1*), taurine uptake (*Slc6a6*/TauT) and taurine consumption via bile-acid conjugation (*Baat*) (Figure 8).

**Figure 8.**
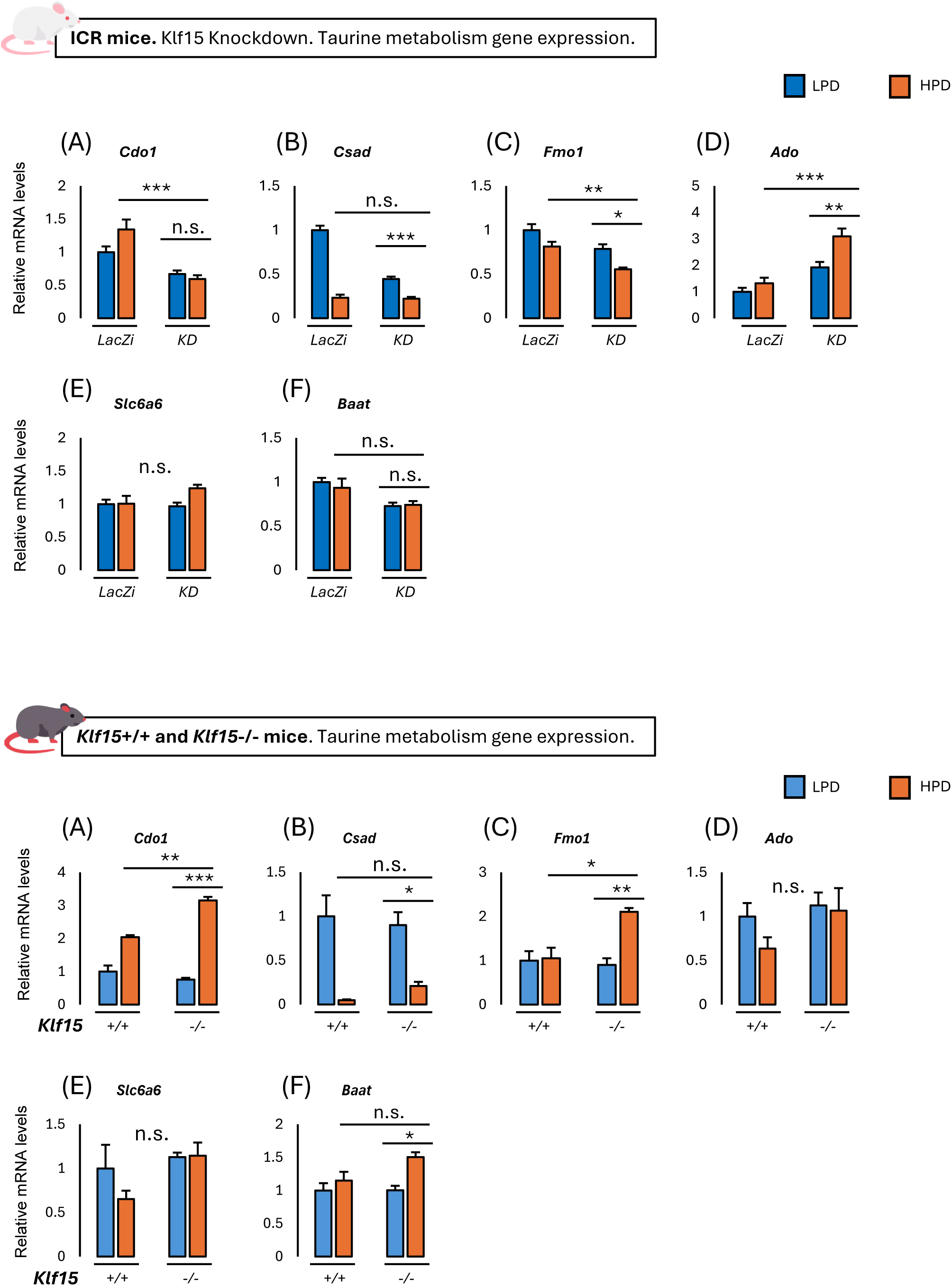
Taurine-biosynthetic gene expression is reprogrammed by Klf15 loss. Relative hepatic mRNA levels of taurine-metabolism genes measured by quantitative RT-PCR (normalized to *36B4*/*Rplp0*) in mice on low-protein diet (LPD, blue) or high-protein diet (HPD, orange). ICR mice injected with control (*LacZi*) or *Klf15*-knockdown (*KD*) adenovirus (Upper panels, A–F) (*n* = 10 per group). *Klf15*⁺/⁺ and *Klf15*⁻/⁻ mice (Lower panels, A–F) (*n* = 3 per group). Genes: **(A)** *Cdo1* (cysteine dioxygenase type 1); **(B)** *Csad* (cysteine sulfinic acid decarboxylase); **(C)** *Fmo1* (flavin-containing monooxygenase 1); **(D)** *Ado* (2-aminoethanethiol [cysteamine] dioxygenase); **(E)** *Slc6a6* (taurine transporter, TauT); **(F)** *Baat* (bile acid-CoA:amino acid *N*-acyltransferase). Data are mean ± SEM (n = 3 to 10 per group). Groups were compared by one-way ANOVA with Tukey HSD post hoc test. \**P* < 0.05, \*\**P* < 0.01, \*\*\**P* < 0.001; n.s., not significant.

In the knockdown model, *Cdo1*, the committed step of taurine synthesis, was HPD-induced in LacZi mice but not in KD mice, and was globally lower after knockdown. *Csad* was strongly suppressed by HPD in both groups (LacZi and KD; group comparison n.s). *Fmo1* was modestly reduced by HPD and further reduced by knockdown. *Ado* was markedly higher in KD mice and HPD-induced within that group. *Slc6a6*/*Taut* and *Baat* were unchanged (n.s.).

In *Klf15*^-/-^ mice, *Cdo1* was HPD-induced in both genotypes and significantly higher in knockout mice under HPD, the opposite direction to that seen with acute knockdown. *Csad* was again HPD-suppressed in both genotypes (genotype comparison n.s.). *Fmo1* was HPD-induced only in *Klf15*^-/-^ mice. *Ado* showed no significant change (n.s.), and *Baat* was modestly HPD-induced in knockout mice only.

The downstream taurine-synthesis genes are variably regulated. The divergence between the two models is pronounced for *Cdo1* and *Ado* and should be interpreted with caution. *Csad* was consistently reduced by HPD in both cohorts and both genotypes. Together these data indicate that HPD-driven hepatic taurine accumulation is not strictly dependent on KLF15 control of the dedicated taurine-synthesis enzymes and are compatible with a model in which the main KLF15 contribution to taurine output is upstream — through *Cth*-dependent provision of cysteine substrate — while the taurine-committed enzymes retain considerable KLF15-independent HPD responsiveness.

### Klf15 is required for full HPD induction of *Cth* and related H₂S-generating genes

Having identified *Cth* (cystathionine γ-lyase) as a direct, HPD-responsive KLF15 target we next asked whether the HPD-driven increase in hepatic hydrogen sulfide (H₂S) is channeled predominantly through *Cth* or is distributed across the broader set of enzymes capable of generating H₂S in the liver. Hepatic mRNA levels of H₂S-metabolizing enzymes were quantified by RT-qPCR (Figure 9). In the knockdown model, *Klf15* itself was reduced and retained a residual HPD response. *Cth* was strongly induced by HPD in control livers, but this induction was significantly attenuated by Klf15 knockdown. *Cbs* was induced by HPD in both groups and was significantly higher in KD livers suggesting a Klf15-independent pattern of expression. *Mto* and *Ast* were induced by HPD and even though their profile was reduced in knockdown group, this effect did not reach significance (n.s.). *3-Mst*, *Cars2* and *Trx* were unchanged by either diet or knockdown (n.s.).

**Figure 9.**
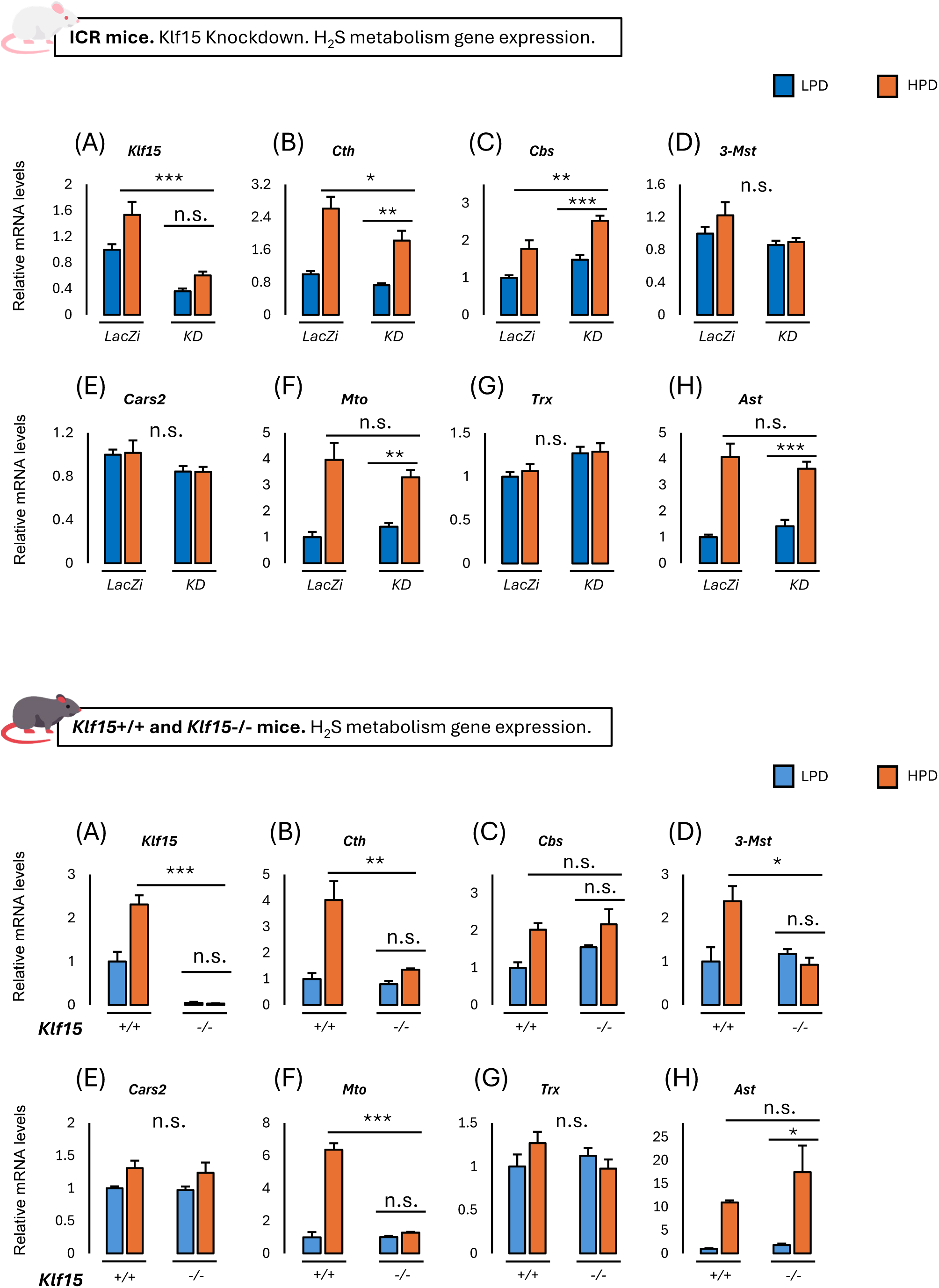
Klf15 is required for full HPD induction of *Cth* and related H₂S-generating genes. Relative hepatic mRNA levels of H₂S-metabolism genes measured by quantitative RT-PCR (normalized to *36B4*/*Rplp0*) in mice maintained on low-protein diet (LPD, blue) or high-protein diet (HPD, orange). ICR mice injected with control (*LacZi*) or *Klf15*-knockdown (*KD*) adenovirus (Upper panels, A–H) (*n* = 10 per group). *Klf15*⁺/⁺ and *Klf15*⁻/⁻ mice (Lower panels, A–H) (*n* = 3 per group). Genes: **(A)** *Klf15* (Krüppel-like factor 15); **(B)** *Cth* (cystathionine γ-lyase); **(C)** *Cbs* (cystathionine β-synthase); **(D)** *3-Mst*/*Mpst* (3-mercaptopyruvate sulfurtransferase); **(E)** *Cars2* (cysteinyl-tRNA synthetase 2); **(F)** *Mto*/*Selenbp1* (methanethiol oxidase / selenium-binding protein 1); **(G)** *Txn1*/*Trx* (thioredoxin); **(H)** *Ast* (aspartate aminotransferase / cysteine aminotransferase activity). Data are mean ± SEM (n = 3 to 10 per group). Groups were compared by one-way ANOVA with Tukey HSD post hoc test. \**P* < 0.05, \*\**P* < 0.01, \*\*\**P* < 0.001; n.s., not significant.

In *Klf15*^-/-^ mice, *Klf15* transcript was undetectable and diet-unresponsive (n.s.). HPD induced *Cth* increase in wild-type livers; this induction was markedly reduced in knockouts. *Mto* showed the same pattern. *3-Mst* was HPD-induced in wild-type but not in knockout livers (n.s.). *Ast* was strongly HPD-induced in both genotypes (n.s. between genotypes), and *Cbs*, *Cars2* and *Trx* were not significantly affected (n.s.).

Across both models, therefore, Klf15 is required for the full transcriptional induction of *Cth* by dietary protein, whereas the responses of *Cbs*, *Cars2* and *Trx* are largely Klf15-independent. Also, we previously established that Ast is expressed independent of Klf15 in response to HPD (9). (primers in Supplementary Table S4). The transcriptional data therefore show a graded effect that parallels the depth of KLF15 loss: partial knockdown blunts *Cth* induction, whereas complete germline deletion abolishes it.

## Discussion

We identify Krüppel-like factor 15 (KLF15) as a direct transcriptional regulator of cystathionine γ-lyase (*Cth*), and we place this interaction at the center of the hepatic sulfur amino-acid response to dietary protein.

In mouse liver, endogenous KLF15 occupies a single discrete region of the proximal *Cth* promoter. Two functional elements within it, 1-1 and 2-2, together account for most of both KLF15-driven and high-protein-diet (HPD)-driven transactivation. When KLF15 is lost, whether by acute adenoviral knockdown in ICR mice or germline deletion in *Klf15*^-/-^ mice, the HPD-induced induction of *Cth* is blunted, the accompanying rise in hepatic hydrogen sulfide (H₂S) is attenuated, and the methionine cycle is incompletely remodeled. The resulting metabolite profile is characteristic: methionine, cystathionine and cystine accumulate while taurine fails to rise fully, a “substrate-up, product-down” signature. Read together, these data favor KLF15 as an amplitude modulator that tunes the magnitude and downstream disposition of dietary sulfur through the CTH-centered pathway, not an obligatory gate without which the response cannot proceed.

The axis established here, HPD → KLF15 → CTH, builds on our earlier finding that *Cth* is a KLF15-dependent, HPD-responsive gene (9). That conclusion had rested only on transcript measurements in *Klf15*-deficient mice, and the responsible cis-regulatory element was never mapped. We can now place it. Public KLF15 ChIP-seq data showed a single enrichment peak immediately upstream of the *Cth* transcription start site, and when this interval was dissected into reporter subfragments, the proximal one (F1) conferred KLF15-responsive luciferase activity in HepG2 cells whereas the distal one (F2) did not. Electrophoretic mobility shift assays traced that activity to two elements: the recombinant KLF15 DNA-binding domain bound 1-1 and 2-2 directly, the overlapping 1-2 and 2-1 elements were dispensable, and mutating both bound elements abolished KLF15-driven reporter activity. Two closely apposed CACCC/GC-rich elements engaged by a single zinc-finger factor is an architecture that fits the known DNA-binding preference of KLF15 (21) and the promoter logic reported for KLF15 at other hepatic targets (29,31). Beyond the reporter, the interaction proved physiologically operative. Endogenous KLF15 occupied both elements in native liver chromatin under HPD, with ChIP-qPCR enrichment comparable to that at the *Gpt* (*Alt*) promoter, a positive KLF15 target locus (24), and none at a gene-desert amplicon. The same elements were needed for the diet response of the intact promoter in vivo: the native promoter and F1, but not F2, were induced by the switch from a low-protein diet (LPD) to HPD, and mutating either element attenuated HPD responsiveness while mutating both largely eliminated it. The effect was graded and selective. Partial knockdown blunted Cth induction whereas germline deletion abolished it, yet *Cbs*, *Cars2*, *Trx* and *Got1* (*Ast*) responded to HPD largely independently of KLF15 (9). KLF15 is thus required for full induction of *Cth* and a small set of co-regulated H₂S-generating transcripts, not as a general amplifier of the sulfur-gene programme.

Most hepatic H₂S is generated through the transsulfuration pathway, and CTH accounts for most of it from cysteine (18). HPD raised hepatic H₂S in control livers, and acute Klf15 knockdown blunted that rise, so KLF15 is needed for the full H₂S response to dietary protein. The rest of the H₂S-generating complement (Cbs, 3-Mst/Mpst, Cars2) behaved largely independently of KLF15 (14,18). The simplest reading is that dietary protein raises endogenous H₂S capacity mainly by amplifying *Cth* transcriptionally, with KLF15 setting the gain. That is a capacity-setting mechanism, working within the existing constraints of substrate supply, compartmentalisation and sulfide clearance, and it differs in kind from delivering a bolus of sulfide with an exogenous donor (16).

Although our experiments do not establish how dietary protein increases hepatic KLF15 activity, this question can be considered in light of existing evidence. Hepatic *Klf15* gene expression is rapidly induced by fasting (23–25), promotes the fasting program of gluconeogenesis and amino acid catabolism, and functions within an insulin-FoxO regulatory circuit in which FoxO1 and FoxO3a directly activate liver-specific *Klf15* transcription (31). Moreover, we previously showed that a high-protein meal increases FoxO binding to the enhancer region of the urea cycle gene *Ass1* (32), suggesting that FoxO may contribute to the hepatic KLF15 response to a high-protein diet (HPD).

The two loss-of-function models diverged: acute knockdown blunted the HPD-induced H₂S response, whereas germline *Klf15*^-/-^ males retained it. This may reflect lifelong upregulation of KLF15-independent sulfide and taurine pathways in germline mutants, which cannot occur after acute adult knockdown. Differences in perturbation scope and completeness, genetic background (ICR vs. C57BL/6J), and age (7–9 weeks vs. 5–6 months) may also alter transsulfuration capacity. Finally, sex-dependent compensation may specifically preserve the male response, as discussed below.

The germline knockout showed a sex-dependent phenotype: the HPD-induced H₂S response was preserved in males but attenuated in females, whereas acute knockdown affected both sexes similarly. Hepatic H₂S production is hormonally regulated (35), and sex hormones modulate transsulfuration and taurine synthesis through CBS and CSAD (36–39). These effects may provide males with greater KLF15-independent compensatory capacity, whereas females may be more vulnerable to impaired CTH function (40). However, because hormones and their receptors were not measured, this explanation remains hypothetical.

We did not report free cysteine on its own; free cysteine was not measured because it readily oxidizes to cystine and is poorly resolved by standard amino-acid analysis (41,42). Because hepatocytes also maintain free cysteine at low levels (10,11,43), cystine was used as a stable surrogate for the cysteine pool produced by CTH, rather than as a separate pathway metabolite.

It might seem paradoxical that cystine accumulates when CTH, a cysteine-producing enzyme, is blunted. Cystine accumulation despite reduced CTH likely reflects sustained cysteine production through abundant cystathionine, CBS, and residual CTH activity (14,18), combined with reduced disposal to H₂S and taurine. Lower H₂S may also favor cysteine oxidation to cystine (17). Thus, the substrate-up, product-down profile supports partial restriction, rather than complete blockade, of CTH-dependent flux.

Hepatic CTH channels dietary sulfur into cysteine and its systemic products, H₂S and taurine (10–12,16–20). Thus, the KLF15-CTH axis may influence systemic sulfur metabolism. Whole-body deletion may additionally recruit extrahepatic, sex-dependent pathways that remain intact after liver-directed knockdown (36–38). The hepatic node described here is one control point in a larger inter-organ sulfur network.

Our findings sharpen the translational rationale for this KLF15-CTH axis in ways tied to the specific results; this axis may have translational relevance because both sulfur amino acid restriction and high-protein feeding increase CTH-dependent H₂S-generating capacity, identifying CTH as a broadly nutrient-responsive control point (44,45). This may be relevant to the age-related decline in taurine and the pathology associated with taurine depletion (46–49). Unlike exogenous H₂S donors, *Cth* induction may enhance endogenous H₂S production within physiological constraints (16). However, the residual response after KLF15 loss and its sex dependence indicate that this axis modulates, rather than fully controls, sulfur metabolism (36–38,40). Because KLF15 is regulated by glucocorticoid receptor and insulin/FoxO signaling and is linked to lipid metabolism, it may provide a node for coordinating lipid and sulfur amino acid metabolism (25,29,31). Further studies are required to establish its therapeutic potential. In summary, KLF15 is a direct transcriptional regulator of *Cth*, and the KLF15-

CTH interaction is a diet-responsive amplifier of hepatic sulfur amino-acid disposal through which dietary protein raises the liver’s endogenous H₂S-generating capacity and shifts the cysteine pool toward taurine. KLF15 is not required for the response, but it sets the response’s amplitude and downstream disposition: a defined, physiologically operative and pharmacologically accessible node that links protein intake to the production of two mediators of therapeutic interest.

## Experimental procedures

### Animals

ICR mice were purchased from Japan SLC, Inc. (Shizuoka, Japan). The experiments using male ICR mice were conducted at age between 7 and 9 weeks old. The *Klf15*^-/-^ (*Klf15*KO) backcrossed into C57BL/6J strain mice was kindly gifted from Prof. Jain MK and genotyped as previously described (28). C57BL/6J *Klf15*^+/+^ (wild type) mice were used as control and KO experiments were performed at age of 5 to 6 months. All animals were housed under a 12-hour light / 12 hour-dark cycle, temperature-controlled environment with free access to food and water. The standard laboratory diet was used at arrival time (Cat#MF; Oriental Yeast, Tokyo, Japan; consisted of 25.5% of energy from protein, 61.5% from carbohydrates, and 13% from fat). High and low protein diets were purchased from Oriental Yeast (Itabashi, Tokyo, Japan) for a 3-day dietary intervention experiments. HPD contained 60 % of energy from casein protein, 27 % from carbohydrate. LPD contained 5% of energy from casein protein, 82 % from carbohydrate. Interventional and arrival diets had the same amount of energy per weight, fat, vitamins and minerals. All animals were anesthetized and euthanized based on the protocol approved by the Tsukuba University and Jichi Medical University Animal Care and Use Committees. To eliminate bias for each experimental environment, they were performed at least twice.

### Measurement of hepatic amino acids

For liver amino acid measurement, 200 mg of liver tissue was homogenized in 2 ml of ice-cold sulfosalicylic acid. Samples were kept on ice for 30 minutes then centrifuged at 5000 rpm for 15 minutes at 4 °C. Supernatant was collected and the pH was adjusted to 2-3 using NaOH. After filtering samples with 0.45-µm filter (Cat# UN203NPUAQU, Mini-UniPrep™, Whatman, Cytiva, Marlborough, MA, USA) at 15,000 rpm for 1 min, amino acid composition was measured with Hitachi-Hightech (LA8080) HPLC machine (32). All stages of this experiment were performed on ice.

### Quantitative reverse transcription PCR (Q-RT PCR)

Up to 100 mg of mouse liver tissue was isolated to extract RNA using Sepasol-RNA I Super G (Cat# 09379-55; Nacalai Tesque, Kyoto, Japan) based on the manufacturer’s instructions. ReverTra Ace qPCR RT Master Mix (Cat# FSQ-201, TOYOBO, Osaka, Japan) was utilized to reverse transcribe 500 ng of extracted RNA in a 5 µL volume to cDNA. KAPA SYBR Fast qPCR Kit (Cat# KK4602, NIPPON Genetics, Tokyo, Japan) was used for Real-time PCR on a QuantStudio™ 5 Real-Time PCR System (Thermo Fisher Scientific, Waltham, MA, USA) and results were quantified by the standard curve method with cDNA as the template. After amplification by PCR, melt curve plot for each sample provided the information for detecting samples containing the product with the correct Tm value. As the internal reference for correction of the gene expression level, *Rplp0* was used for all samples. Primer sets are shown in Table S4.

### Cell culture

HEK293 human embryonic kidney cells (RRID: CVCL_0045) and HepG2 human hepatoma cells (RRID: CVCL_0027) were obtained from ATCC (American Type Culture Collection, Manassas, VA, USA), and cultured in DMEM containing 25 mM glucose, 100 U/mL penicillin, and 100 μg/mL streptomycin sulfate supplemented with 10% FBS or 10% CCS. All cell lines have been authenticated by our facility’s authentication protocol within the last 3 years. Experiments were performed with mycoplasma-free cells.

### In vivo imaging of luciferase activity

In vivo imaging was performed as described previously (29,33,34). After 5 days of the adenovirus transduction, animals were fed LPD or HPD for a duration of 3 days. At each condition, D-Luciferin potassium salt (Cat# 126-05116, Wako Chemicals, Tokyo, Japan) dissolved in PBS at a concentration of 7.5 mg/mL was intraperitoneally injected at a dose of 10 mL/kg into mice. The luminescence in the liver was captured using an IVIS™ Imaging System (PerkinElmer, Waltham, MA, USA) and Living Image™ software (PerkinElmer) was used to quantify relative photon emissions over the liver region. Two paired data from the same animal on the different nutritional conditions (i.e., LPD or HPD) was continuously obtained and the ratio between the two quantities was used to cancel the variations in hepatic transduction efficiencies. Results less than 1 x 105 counts/min (2.44 x 106 photons) on both the conditions were not adopted due to inadequate detection accuracy.

### Chromatin immunoprecipitation (ChIP) assay

Mouse liver tissue was subjected to Chromatin immunoprecipitation (ChIP) assays as described previously (31). 100 mg of liver tissue samples from the HPD-fed mice were minced in 1 mL of PBS and cross-linked in 1.5% formaldehyde for 15 min at room temperature as fixation stage. Minced samples were homogenized and to fragment the DNA, Bioruptor2 (Sonicbio, Kanagawa, Japan) was used for sonication. The mixture was centrifuged and supernatant was diluted to 6 mL with dilution buffer (50 mM Tris-HCl at pH 8.0, 167 mM NaCl, 1 mM EDTA, 1.1% Triton X-100, and 0.1% sodium deoxycholate). 1 mL of total volume was used for immunoprecipitation with 1 µg of anti-Klf15 (Cat# sc-271675, Santa Cruz Biotechnology) or control IgG (Cat# sc-2025, Santa Cruz Biotechnology) bound by 30 µL of Dynabeads magnetic beads (Thermo Fisher Scientific, Waltham, MA, USA) and rotated overnight at 4 °C. Fifty microlitres of total volume was used for input sample. The complexes were washed with low-salt wash buffer (50 mM Tris-HCl at pH 8.0, 150 mM NaCl, 1 mM EDTA, 1% Triton X-100, 0.1% SDS, and 0.1% sodium deoxycholate), high-salt wash buffer (50 mM Tris-HCl at pH 8.0, 500 mM NaCl, 1 mM EDTA, 1% Triton X-100, 0.1% SDS, and 0.1% sodium deoxycholate), LiCl wash buffer (10 mM Tris-HCl at pH 8.0, 0.25 M LiCl, 1 mM EDTA, 0.5% NP40, and 0.5% sodium deoxycholate), and TE buffer. Elution buffer (1% SDS, 0.1 M NaHCO3) was used to incubate DNA-protein complex at room temperature for 15 min. Reverse crosslinking step occurred overnight at 65 °C through incubation with 200 mM NaCl. 200 μg / mL proteinase K treatment was applied on DNA-protein complex, and chromatin DNA was purified with phenol-chloroform. Samples were eluted in TE buffer, and were used for Q-PCR analysis. The input sample was also subjected to the same procedure from the reverse-crosslinking step. Q-PCR was performed using the same method as Q-RT PCR and quantified by standard curve method with input DNA samples. The primer sets are listed in Table S3.

### Preparation and transduction of recombinant adenoviruses

Construction of *Cth* promoter firefly luciferase reporter plasmid is briefly described in (9). Figure S1 shows UCSC Genome Browser tracks for H3K4me1, H3K27ac, and H3K9ac. The smaller fragments of *Cth* promoter region were amplified by PCR using mouse genomic DNA as template and inserted into multiple cloning site on the pENTR4-Luc Gateway entry vector linked to firefly luciferase reporter (Cth-Luc) as previously described (50). Mutated Cth-luc plasmids were generated using PrimeSTAR Mutagenesis Basal Kit (Cat# R046A, TaKaRa Bio, Tokyo, Japan) based on PCR. The primer sets for PCR and brief methods are listed in Table S1. Adenoviral constructs were generated by homologous recombination between the entry vector and the pAd promoterless vector (Thermo Fisher Scientific, Waltham, MA, USA) according to the manufacturer’s protocol.

Adenovirus encoding LacZi-specific shRNA for RNAi (LacZi) and Klf15-specific shRNA for RNAi (Klf15i) were described previously (29). Briefly, LacZi or Klf15-specific shRNA construct inserted in pENTR/U6 the entry vector was homologously recombined with the adenovirus vector. Recombinant adenoviruses were propagated in HEK293 cells and purified by CsCl gradient centrifugation as described previously (51). Solution 1 (5.30 g of CsCl in 8.7 mL of 10 mM Tris-HCl at pH 7.9) and Solution 2 (2.68 g of CsCl in 9.2 mL of 10 mM Tris-HCl at pH 7.9) were prepared, and first 1.02 mL of solution 1, then 0.78 mL solution 2 were layered in 3.5 ml polypropylene Quick-Seal centrifuge tube (Beckman Coulter). The cell lysate containing recombinant adenovirus was gently overlaid and centrifuged for 3 hours at 89,000 rpm at 4 °C. The tube was punctured at the bottom with a 21-gauge needle, and fraction containing adenovirus was collected. To further purify recombinant adenovirus, the collected sample was fractionated on a PD-10 desalting columns containing Sephadex G-25 resin (GE Healthcare) by adding phosphate-buffered saline (PBS) with 1 mM MgCl_2_. For animal experiments, adenoviruses were injected intravenously into mice via the jugular vein at the following doses: for all Cth-Luc (F1, F2, F1(1–1) mut, F1(2–2) mut and F1(Double) mut), 1.0 × 10⁸ PFU and for LacZi/Klf15i, 2.0 × 10^9^ PFU. 1000 optical particles of adenovirus were calculated as 1 PFU.

### Luciferase assay and TFEL scan genome-wide transcription factor screening

For luciferase assay cells were seeded in 48-well plate to 20% confluence and transfected using Lipofectamine 3000 reagent (Cat# L3000-015, Thermo Fisher Scientific, Waltham, MA, USA) according to the manufacturer’s protocol as described previously (30,31). Cells were transfected with Klf15 expression plasmids (31), *Cth*-promoter firefly luciferase reporter plasmids and Renilla luciferase reporter plasmids (pRL-SV40; Promega, Madison, WI, USA) after adjusting total amounts of transfected DNA with empty plasmid. After 48 h, cells were lysed with 100 μL of Reporter Lysis Buffer (Cat# E397A, Promega, Madison, WI, USA) and centrifuged at 15,000 rpm for 15 min at 4 °C. The supernatant was mixed with a luminometer with a firefly luciferase assay reagent (Cat# PGL5500, Pikkagene, Toyo Bnet bio, Tokyo, Japan) and the firefly luciferase activity was measured using a Wallac ARVO SX 1420 luminometer (PerkinElmer, Shelton, CT, USA). Renilla luciferase activity was measured with Renilla Luciferase Assay System (Cat# E2820, Promega, Madison, WI, USA) according to the manufacturer’s protocol and was used to normalize transfection efficiencies.

### Electrophoretic mobility shift assay (EMSA)

EMSA was performed as described previously (52). In brief, DNA probes were prepared by annealing two complementary oligonucleotides, labeled with [α-³²P] dCTP by filling in the 5′-overhangs with Klenow DNA polymerase, and purified on Sephadex G-50 columns. EMSA probes are listed in Table S2. A DNA fragment encoding the amino acid sequence containing the mouse KLF15 DNA-binding domain, comprising the three C2H2 zinc-finger motifs, was inserted into the designated plasmid to prepare a GST-fusion protein (GST-KLF15-DBD). GST and GST-KLF15-DBD were expressed in *E. coli* and purified on glutathione-Sepharose (Amersham BioSciences, Buckinghamshire, UK) using standard methods as described previously (29). The labeled DNA probes were incubated with GST (200 ng, as a negative control) or GST-KLF15-DBD (200 ng) in binding buffer (10 mM Tris-HCl (pH7.5), 200 mM KCl, 0.5 mM EDTA, 1 mM MgCl_2_, 1 mM DTT, 5% glycerol, 5 mM ZnCl_2_, 400 ng poly (dI-dC)) for 30 min on ice. For competition assays, a 100-fold molar excess of unlabeled wild-type or mutant oligonucleotide was added to the reaction before addition of the labeled probe. The DNA-protein complexes were analyzed on 4.6% polyacrylamide gels in TBE buffer and visualized by BAS imaging plate on a BAS2000 Bio-Imaging Analyzer.

### H_2_S measurement

Free and acid-labile hydrogen sulfide (H₂S) in liver was quantified spectrophotometrically by the methylene blue method (53). The protocol as modified for murine tissue was described previously (54). Immediately after sacrifice, livers were excised and homogenized in ice-cold 10 mM NaOH at a tissue-to-buffer ratio of 1:5. All subsequent steps were performed on ice. Then, 50% trichloroacetic acid (TCA) was added to homogenate samples making a 10% final concentrate followed by vigorous mixing and centrifugation at 6,000 rpm for 5 min. A 1.0 mL aliquot of the supernatant was transferred to a sealed tube containing 100 µL of 20 mM N,N-dimethyl-*p*-phenylenediamine sulfate in 7.2 M hydrochloric acid (HCl), after which 100 µL of 30 mM iron (III) chloride (FeCl₃) in 1.2 M hydrochloric acid (HCl) was added to drive methylene blue formation. Reactions were incubated for 20 min in the dark, clarified with 800 µL of chloroform, and shaken for 1 min. A blank containing all assay components except tissue homogenate was processed in parallel through the complete procedure, and its absorbance was subtracted from all sample readings. Each sample was loaded in triplicate and read immediately at 600 nm, the wavelength available on the instrument, with the mean of the three wells taken as the value for that animal. Absorbance was calibrated against a sodium sulfide (Na2S) standard curve prepared and read under identical conditions.

### Statistical analyses

Statistical analyses were performed using IBM SPSS Statistics 31.0.0 (IBM Corp., Armonk, NY, USA), with individual mice as the experimental unit for all in vivo experiments. Data are presented as mean ± SEM with individual values. Normality and homogeneity of variance were assessed using the Shapiro-Wilk test with Q-Q plots and Levene’s test, respectively; variances were homogeneous in all reported comparisons. Hepatic metabolites, including H₂S, and gene expression were compared among diet- and genotype- or treatment-defined subgroups by one-way ANOVA with Tukey’s HSD test. Sex comparisons (Supplementary Figure S3) additionally included sex in subgroup definitions. Non-normal data were analyzed using the Kruskal-Wallis test followed by Dunn’s test with Bonferroni correction. Promoter-reporter activity in HepG2 cells (Figure 1) and ChIP-qPCR enrichment relative to IgG (Figure 3) were compared using unpaired Student’s t-tests. Paired before-and-after hepatic photon flux measurements (Figures 2C and 2F) were compared using the Wilcoxon signed-rank test. Activity relative to the corresponding LPD group was compared among constructs using the Kruskal-Wallis test with Bonferroni correction (Figure 2B) or one-way ANOVA with Tukey’s HSD test (Figure 2E), depending on normality. Animals with hepatic luminescence below 1 × 10⁵ counts/min under both diets were excluded before analysis, resulting in group sizes of *n* = 8–14. All tests were two-tailed, with *P* < 0.05 considered significant (*P* < 0.05, \**P* < 0.01, and \*\**P* < 0.001).

## Acknowledgements

We thank Prof. Mukesh Jain (Case Western Reserve University) for kindly providing the KLF15 knockout mice.

This work was supported by MEXT/JSPS KAKENHI Grant Numbers 23116006 (Grant-in-Aid for Scientific Research on Innovative Areas: Crosstalk of transcriptional control and energy pathways by hub metabolites), 23K24760 and 15H03092 (Grant-in-Aid for Scientific Research (B)), and 24K22110 (Grant-in-Aid for Challenging Research (Exploratory)) (to N. Yahagi). This research was also supported by AMED under Grant Numbers JP23gm1710008 (AMED-CREST) and JP23rea522010 (Healthcare Social Implementation Infrastructure Development Project) (to N. Yahagi). This research was also supported by MEXT/JSPS KAKENHI Grant Number 23K13900 (Grant-in-Aid for Early-Career Scientists) (to Z. Saber).

## Author contributions

Z.S. and N.Y. conceived the experiments. Z.S. performed the experiments and analyzed the data together with N.Y. Z.S. and N.Y. co-wrote the manuscript. All authors discussed the results and commented on the manuscript.

## Abbreviations

CTH: Cystathionine gamma-lyase
HPD: high-protein diet
KD: knockdown
KLF15: Krüppel-like factor 15
KO: knockout
LPD: low-protein diet

