## Supporting Information for "KLF15 regulates sulfur amino acid metabolism through Cystathionine gamma-lyase"

(A)

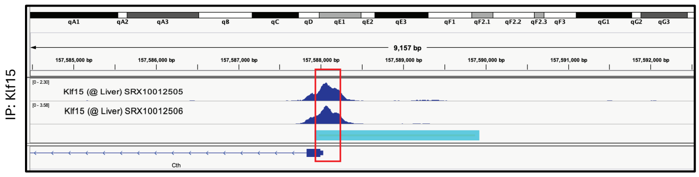


(B)

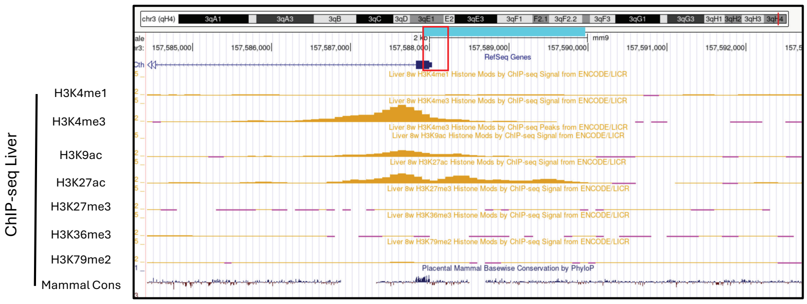


(C) Fragment 1 sequence

TGAGTGCTGCACCTCAGTGCTGAGGTGTCCCACGTGGGCACTGCTCTGTGCCACTGGGAGGCGGGGCAGGAA

CGATCGGGGCAACACCTCCACTGCCGACCAATAAGCAGGGCCGCTGCCTAGGGACCAGCGATGATTGGTTGC

GTCGGCCCCTCCCCACCCTGGATATAAGCGCCAAAGTCCAGAAGGTTTTAGCCAAAGCAACACCTCGCACTC

(D) JASPAR 2022 scores for predicted binding sites in fragment 1

| Matrix ID | Name | Score | Relative score | Strand | Predicted sequence |
|---|---|---|---|---|---|
| MA1513.1 | MA1513.1.KLF15 | 9.303394 | 0.876312448767364 | − | GCCCCGCCTCC |
| MA1513.1 | MA1513.1.KLF15 | 6.9307337 | 0.8420739038510674 | + | GCCCCTCCCCA |
| MA1513.1 | MA1513.1.KLF15 | 4.973971 | 0.8138369492452394 | − | TTCCTGCCCCG |
| MA1513.1 | MA1513.1.KLF15 | 4.855002 | 0.8121201743549323 | + | TCCCCACCCTG |

**Figure S1**

**(A)** Publicly available Klf15 chromatin immunoprecipitation–sequencing (ChIP-seq) data from adult mouse liver (accessions SRX10012505 and SRX10012506). Blue marker represents the *Cth* promoter whole fragment. **(B)** Modified Histones binding regions on *Cth* promoter from ENCODE ChIP-seq database (Liver 8w by ChIP-seq Signal from ENCODE/LICR) on Genome browser. These peaks indicate promoter/enhancer regions. Mammalian conservation regions are shown below ChIP-seq peaks data. Blue marker represents the *Cth* promoter whole fragment. **(C)** Mouse genomic sequence of fragment 1. In red are the four predicted Klf15 binding sites overlapping as two pairs. **(D)** *In silico* scanning of the 216-bp F1 sequence against the JASPAR 2022 CORE vertebrate matrix for KLF15 (MA1513.1) was used to predict Klf15 binding sites. (https://jaspar2022.elixir.no/)

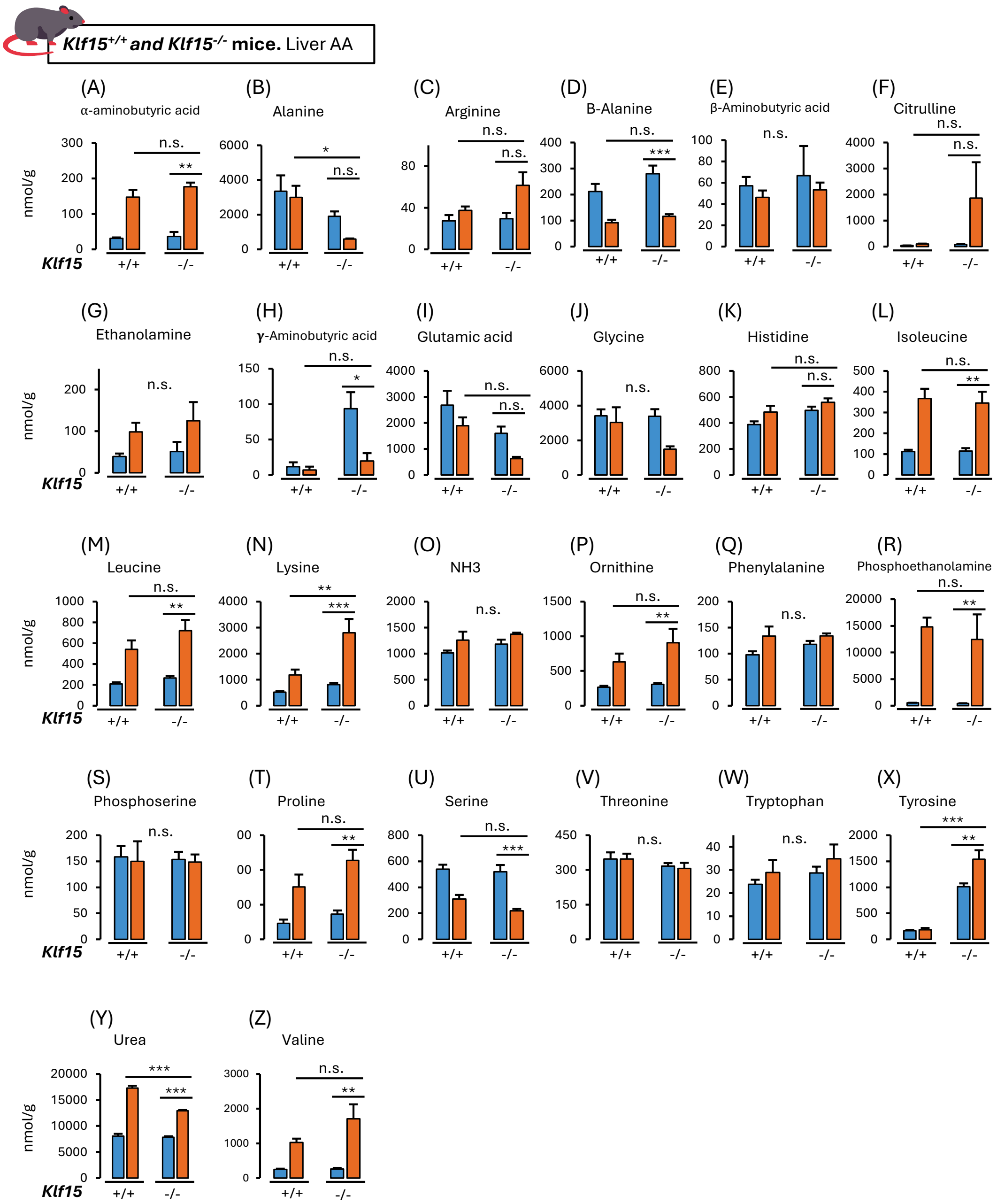


### Figure S2

Amino acid measurements were made in germline *Klf15*$^{-/-}$ mice and *Klf15*$^{+/+}$ littermates. Hepatic concentrations (nmol/g tissue) of free amino acids and related metabolites are shown under LPD (blue) and HPD (orange). **(A–Z)** panels show the 26 amino acids aside from methionine, cystathionine, cystine, taurine. $n$ = 6 per group and data are mean ± SEM. Group comparisons were assessed by ANOVA with Tukey's HSD post hoc test for normally distributed data and by the Kruskal-Wallis test with Bonferroni correction for non-normally distributed data; $^{*}P < 0.05$, $^{**}P < 0.01$, $^{***}P < 0.001$, n.s. = not significant.

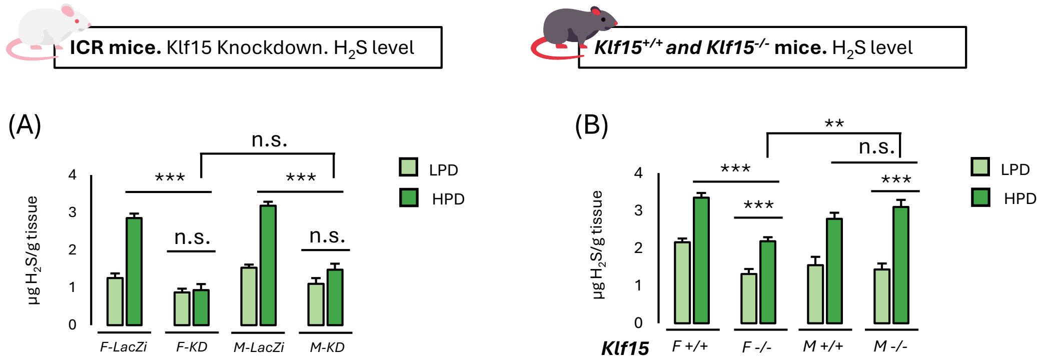


**Figure S3**

**(A)** Hepatic $H_2S$ content (μg $H_2S$/g tissue) in female (F) and male (M) ICR mice after adenoviral control (F-*LacZi*, M-*LacZi*) or *Klf15* knockdown (F-*KD*, M-*KD*), under LPD and HPD ($n$ = 6 per group). **(B)** Hepatic $H_2S$ content in female and male *Klf15*$^{+/+}$ (F +/+, M +/+) and *Klf15*$^{-/-}$ (F −/−, M −/−) mice under LPD and HPD ($n$ = 6 per group). Data are mean ± SEM. Diet and genotype/treatment effects within each sex were analyzed by one-way ANOVA with Tukey's HSD post hoc test (normally distributed data) or the Kruskal-Wallis test with Bonferroni correction (non-normally distributed data); sex was examined by assigning animals to subgroups defined by diet and sex and comparing through one-way ANOVA followed by Tukey's HSD post hoc test. Analyses were performed in SPSS version 31.0.0 with α = 0.05. *$P$ < 0.05, **$P$ < 0.01, ***$P$ < 0.001; n.s., not significant. Panels A and B are independent experiments, separate from the male-only cohort in Figure 6 and assayed with a refined $H_2S$ protocol; absolute values are therefore not directly comparable across figures.

**Table S1. List of primer sets used for Q-RT PCR.**

| *Cth*-Ptomoter-Luc | Sequence for PCR |
|---|---|
| Full | 5′-AGGAATCCATGGGTTAGCAGTG-3′<br>5′-CAAAGCAACACCTCGCACTC-3' |
| F1 | 5′-TGAGTGCTGCACCTCAGT-3′<br>5′-CAAAGCAACACCTCGCACTC-3′ |
| F2 | 5′-AGCGATGATTGGTTGCGT-3′<br>5′-CAAAGCAACACCTCGCACTC-3′ |
| F1 (1-1) Mutated | 5′-TGGGAGGATCCGCAGGAACGATCGGGGCA-3′<br>5′-CTGCGGATCCTCCCAGTGGCACAGAGCA-3′ |
| F1 (2-2) Mutated | 5′-CGGCGGATCCCCACCCTGGATATAAGCG-3′<br>5′-GGTGGGGATCCGCCGACGCAACCAATCAT-3′ |

**Table S2. List of primer sets used for ChIP assay.**

| Probe | Sequence |
|---|---|
| Site 1 WT | 5′-GTGCCACTGGGAGGCGGGGCAGGAACGATCGGGG-3′<br>5′-GTTGCCCCGATCGTTCCTGCCCCGCCTCCCAGTGG-3′ |
| Site 2 WT | 5′-GGTTGCGTCGGCCCCTCCCCACCCTGGATATAAG-3′<br>5′-GCGCTTATATCCAGGGTGGGGAGGGGCCGACGCAA-3′ |
| Site 1(1-1) mut | 5′-GTGCCACTGGGAGGATCCGCAGGAACGATCGGGG-3′<br>5′-GTTGCCCCGATCGTTCCTGCGGATCCTCCCAGTGG-3′ |
| Site 1(1-2) mut | 5′-GTGCCACTGGGAGGCGGGATCCGAACGATCGGGG-3′<br>5′-GTTGCCCCGATCGTTCGGATCCCGCCTCCCAGTGG-3′ |
| Site 2(2-1) mut | 5′-GGTTGCGTCGGCCCCTCCCGGATCCGGATATAAG-3′<br>5′-GCGCTTATATCCGGATCCGGGAGGGGCCGACGCAA-3′ |
| Site 2(2-2) mut | 5′-GGTTGCGTCGGCGGATCCCCACCCTGGATATAAG-3′<br>5′-GCGCTTATATCCAGGGTGGGGATCCGCCGACGCAA-3′ |

**Table S3. List of primer sets used for ChIP assay.**

| Gene (mouse) | Sequences for ChIP Q-PCR primers |
|---|---|
| *Cth 1-1* | 5′-CTGAGTGCTGCACCTCAGT-3′<br>5′-ACGCAACCAATCATCGCTGGT-3′ |
| *Cth 2-2* | 5′-ACTGCCGACCAATAAGCAG-3′<br>5′-GAGTGCGAGGTGTTGCTTTGGCT-3′ |
| *Gpt* (*Alt*) | 5′-AACTAGCTGTCCCGTCTCCA-3′<br>5′-CTCTGATGAGCCACTGCAAG-3′ |
| *Negative Control* | 5′-GGGCACATAATGCCTTTCTGA-3′<br>5′-CTGCCTCTGGAGTGATGGGATTA-3′ |

**Table S4. List of primer sets used for Q-RT PCR.**

| Gene (mouse) | Sequences for Q-RT PCR primers |
|---|---|
| *3-Mst / Mpst* | 5′-CTCACAGCCGCTGAAGTTACTG-3′<br>5′-GTGGCGTTAGGCAGCATGT-3′ |
| *Ado* | 5′-CACCTACATGCACATCTACG-3′<br>5′-GTACAGCACCTTGAGCATAC-3′ |
| *Ahcy* | 5′-GGAGACTGCTGTTCTCATTGAG-3′<br>5′-TGCATGGTCCTGAGTAGAGAA-3′ |
| *Baat* | 5′-CTCTGGAAAGGTGGTATGTG-3′<br>5′-GCACCTCCAAACAAGTCAAT-3′ |
| *Bhmt* | 5′-GATTGGAGATGGAGGATTTG-3′<br>5′-GTCTGCATGACGTTCGAT-3′ |
| *Cars2* | 5′-AGCTGCTCTAGCCAATGACT-3′<br>5′-GAGTTCATCTACCACACAGC-3′ |
| *Cbs* | 5′-TCTGGACCAGTACCGCAATG-3′<br>5′-CTCATAGGCTGTTTGCTCCG-3′ |
| *Cdo1* | 5′-GACCTCATCCGCATCTTACA-3′<br>5′-CCCAGCACAGAATCATCAGA-3′ |
| *Csad* | 5′-TACAGTGTCAAGACTGGTCACC-3′<br>5′-ACTGGCTAGTGTTGAGGCTC-3′ |
| *Fmo1* | 5′-CTGTGGTTTCTAACAGCAGC-3′<br>5′-CTTCCCACTGTCCAGAGACA-3′ |
| *Gnmt* | 5′-TTCTGACCTCATCCAGAGTG-3′<br>5′-AGTTGGCTTCTTCAATGACC-3′ |
| *Klf15* | 5′-GCGAGAAGCCCTTTGCCT-3′<br>5′-GCTTCACACCCGAGTGAGAT-3′ |
| *Mat1a* | 5′-AAGTGAAGAGGGAGCCTTCA-3′<br>5′-GTCTTGCACACTGTCTCACA-3′ |
| *Mthfr* | 5′-CACCCTCTATCAGCAGGAAT-3′<br>5′-GGGTGGAACATCTTGAACTG-3′ |
| *Mto / Selenbp1* | 5′-TGTTAGAGGAGGCTCTGTGC-3′<br>5′-GGTAACGAAGCTCGTGAGCC-3′ |

| | |
|---|---|
| *Mtr / Ms* | 5′-TACTAGTATCGCCCAGGCTG-3′<br>5′-ACCTCTTGACTCCTGTCTGC-3′ |
| *Shmt1* | 5′-GCTGATTGCCTCGGAGAATT-3′<br>5′-TGGTAGGCCTGTAATGCACG-3′ |
| *Shmt2* | 5′-CTCATCGCCTCAGAGAACTT-3′<br>5′-CTGAGTATGGCTGCACATTGAC-3′ |
| *Slc6a6 / Taut* | 5′-ATAGGCCAGTACACATCAGA-3′<br>5′-CTGTGGTGTGTTCCAGCTAT-3′ |
| *Txn1 / Trx* | 5′-TCTCTGCTACGTGGTGTGGA-3′<br>5′-GTTAGCACCGGAGAACTCCC-3′ |
| *36B4 / Rplp0* | 5′-GAAGACAGGGCGACCTGGAA-3′<br>5′-TTGTCTGCTCCCACAATGAAGC-3′ |
